# Genomic Epidemiology of SARS-CoV-2 in Norfolk, UK, March 2020 – December 2022

**DOI:** 10.1101/2024.09.05.611382

**Authors:** Eleanor H. Hayles, Andrew J. Page, Robert A. Kingsley, Javier Guitian, The COVID-19 Genomics UK Consortium, Gemma C. Langridge

**Affiliations:** Quadram Institute Bioscience, Norwich Research Park, Norwich, UK; Norwich Medical School, University of East Anglia, Norwich Research Park, Norwich, UK; Theiagen Genomics, 1745 Shea Center Drive, Highlands Ranch, Colorado, USA; School of Biological Sciences, University of East Anglia, Norwich Research Park, Norwich, UK; Department of Pathobiology and Population Sciences, The Royal Veterinary College, London, UK

**Author notes:** **Corresponding author:** Gemma C. Langridge.

**Keywords:** SARS-CoV-2, genomic epidemiology, variant of concern, Norfolk, COG-UK

## Abstract

**Background:** In the UK, the COVID-19 Genomics UK Consortium (COG-UK) established a real time national genomic surveillance system during the COVID-19 pandemic, producing centralised data for monitoring SARS-CoV-2. As a COG-UK partner, Quadram Institute Bioscience (QIB) in Norfolk sequenced over 87,000 SARS-CoV-2 genomes, contributing to the region becoming densely sequenced. Retrospective analysis of SARS-CoV-2 lineage dynamics in this region may contribute to preparedness for future pandemics.

**Methods:** 29,406 SARS-CoV-2 whole genome sequences and corresponding metadata from Norfolk were extracted from the COG-UK dataset, sampled between March 2020 and December 2022, representing 9.9% of regional COVID-19 cases. Sequences were lineage typed using Pangolin, and subsequent lineage analysis carried out in R using RStudio and related packages, including graphical analysis using ggplot2.

**Results:** 401 global lineages were identified, with 69.8% appearing more than once and 31.2% over ten times. Temporal clustering identified six lineage communities based on first lineage emergence. Alpha, Delta, and Omicron variants of concern (VOC) accounted for 8.6%, 34.9% and 48.5% of sequences respectively. These formed four regional epidemic waves alongside the remaining lineages which appeared in the early pandemic prior to VOC designation and were termed ‘pre-VOC’ lineages. Regional comparison highlighted variability in VOC epidemic wave dates dependent on location.

**Conclusion:** This study is the first to assess SARS-CoV-2 diversity in Norfolk across a large timescale within the COVID-19 pandemic. SARS-CoV-2 was both highly diverse and dynamic throughout the Norfolk region between March 2020 – December 2022, with a strong VOC presence within the latter two thirds of the study period. The study also displays the utility of incorporating genomic epidemiological methods into pandemic response.

**Data summary:** The COG-UK collection of SARS-CoV-2 sequences and metadata are available for public download on their archive website under the ‘Latest sequence data’ heading <https://webarchive.nationalarchives.gov.uk/ukgwa/20230507102210/https://www.cogconsortium.uk/priority-areas/data-linkage-analysis/public-data-analysis/>. Sequence names for all sequences used from this dataset alongside GISAID accession numbers where present are available in **Supplementary Table 1**.

**Impact statement:** We extracted 29,406 regional Norfolk based SARS-CoV-2 sequences from the COG-UK SARS-CoV-2 dataset and revealed significant regional diversity and dynamic emergence of variant of concern (VOC) epidemic waves – spanning Alpha, Delta and Omicron lineages. We also applied statistical modelling to complement genomic methodology, with temporal clustering of significant first lineage emergences chronologically matching VOC waves and subwaves. The study highlights the importance of integration of genomic epidemiology into public health strategies for pandemic response, and the utility of using this data for retrospective research.

## Introduction

Since its emergence and establishment as a global pathogen, Severe Acute Respiratory Syndrome Coronavirus 2 (SARS-CoV-2) has posed one of the biggest public health threats in the last century (1,2). SARS-CoV-2 lineages describe further subdivision between closely related viruses within the species, with Pango classification widely adopted for nomenclature, although further nomenclature systems exist (3–5). Variants of concern (VOC) are specific lineages, or groups of related sublineages which have an increased epidemic potential (such as via enhanced immune evasion or viral entry), caused by the presence of a single or combination of mutations (6,7). VOCs were designated at the global level by the World Health Organisation from late 2020, with a Greek letter identifier assigned to each for ease of public communication from May 2021 (8,9).

An initial period of viral evolutionary stasis was observed from emergence to late 2020 until VOCs emerged and outcompeted existing circulating strains (10,11). There have been five main VOCs in the UK – Alpha (B.1.1.7), Beta (B.1.351), Delta (B.1.617.2 and AY sublineages), Gamma (P.1) and Omicron (B.1.1.529, alongside BA and related sublineages) (12,13). Alpha was first detected in the UK in September 2020 and designated a VOC in December 2020 (13,14). Delta was first detected in December 2020, and was designated a VOC in May 2021, followed by Omicron, which was first recorded in the UK and designated a VOC in November 2021 (15–17). Beta and Gamma VOCs were less prominent than other VOCs, and were first recorded in the UK in December 2020 and February 2021 respectively (13,18).

Whole genome sequencing (WGS) of SARS-CoV-2 has played a pivotal role in global pandemic response, including initial identification of the causative agent, and tracking diversity in near real time since emergence (19–22). Furthermore, the wealth of data produced now provides the opportunity for in-depth, retrospective analyses that have the potential to contribute to preparedness for future pandemics (23,24). Centralised national sequencing capacity for SARS-CoV-2 was established in the UK by the COVID-19 Genomics UK consortium (COG-UK), a partnership between academic institutions, public health bodies and the National Health Service (NHS) (25,26). COG-UK facilitated nationwide coverage of SARS-CoV-2 WGS and genomic analyses of samples for real time surveillance and monitoring, contributing to the UK’s public health response (27). Quadram Institute Bioscience (QIB) in Norfolk was an academic partner, sequencing over 87,000 SARS-CoV-2 genomes and contributing to Norfolk becoming one of the most densely sequenced regions in the UK (28).

Here we characterise the lineage dynamics and VOC diversity of SARS-CoV-2 in Norfolk, UK across the core timeframe of the COVID-19 pandemic, between March 2020 and December 2022, using the COG-UK dataset. We apply statistical modelling methods to further characterise patterns of lineage emergence. By characterising viral evolution and diversity through the main VOC epidemic waves of the COVID-19 pandemic in one well sequenced region, our insights into its diversity provide a retrospective view of a pandemic virus.

## Materials & Methods

### Norfolk data filtering from the COG-UK dataset

Both sequence data and metadata were produced as part of COG-UK pipelines as previously described (27). Metadata was accessed through the COG-UK Cloud Infrastructure for Microbial Bioinformatics (CLIMB) server on 12/01/2023 (29). 5,284,065 entries were present in the UK dataset at the time of metadata download, with 29,406 samples from Norfolk. UK data were filtered for Norfolk using grep() pattern searching in the command line. All metadata entries were dated between 12/03/2020 and 30/12/2022. Sequence data were downloaded from the public COG-UK dataset on 15/06/2023. 3,392,463 sequences were present in this dataset and was filtered to contain the 29,406 Norfolk samples using Seqfu (v1.17.1) (30).

### Norfolk metadata description

Sequence metadata included the sequence name, GISAID accession numbers, sample date, epidemiological week of sample collection, country, and county of sample location (31). Patient specific metadata included age, sex and location recorded to the UK county level. Location was determined via the patients registered address on their NHS record, rather than location of sample. Varying completeness was present across the dataset for each value. QIB involvement with COG-UK allowed access to a higher resolution of metadata – location information beyond country of location, or any patient information is not available publicly.

### Case/Sequence analyses

New daily case numbers for Norfolk by specimen date were downloaded from the UK Government COVID-19 Dashboard (http://coronavirus.data.gov.uk/details/downloads), with Upper Tier Local Authority and Norfolk specified as ‘area type’ and ‘area name’ respectively, and ‘newCasesBySpecimenDate’ specified under ‘metrics’ on 24/05/23. The output was restricted to only include cases from March 2020 to December 2022 to match sequencing dates. Further analyses were carried out using R (v4.3.2) in RStudio (v2023.12.1+402). Dates were converted from daily data to weekly data using aggregate() within the base stats package (v3.6.2) and group_by() and count(), part of the package dplyr (v1.1.2) (32–34). Graphical visualisation undertaken using ggplot2 (v3.4.2) (35). Dates of key UK intervention measures including lockdowns, testing and vaccine availability were overlaid onto plots as appropriate.

### Lineage diversity

Pangolin (v4.3) was used to determine the lineage of each sequence and produce a sequence alignment, with Scorpio (v3.17) assigning a VOC identity to a sequence (31). Analyses were carried out in R in RStudio, using Tidyverse (v2.0.0) packages with graphical visualisation carried out in ggplot2 (33–36). Restriction metrics for determining a lineage list for further analysis were determined using percentile data for the number of times a lineage appeared overall using percentile() within the base stats package (33). Lineages appearing within the top 10^th^ percentile were used in further lineage analysis, creating the focused dataset. Descriptive statistics (including mean, standard deviation and standard error) were calculated using describe() within psych (v2.3.6) (37). Shannon’s Diversity Index (H) was calculated using the vegan package (v2.6-4) (38).

### Phylogenetics

An initial tree was created using RapidNJ (v2.2.6) using the alignment produced as part of Pangolin as the input (31,39). A custom perl script was applied to the output to remove single quotations around sequence names to ensure the format matched the alignment input. IQTree2 (v2.3.2) was used with the following flags: -s (input) as the alignment produced from Pangolin, -t (initial tree) as the tree produced from RapidNJ, -T (number of threads) as 32, -m (chosen model) as HKY+G and –keep-ident to keep identical sequences within the tree. The tree was rooted by the earliest entry in the Norfolk COG-UK dataset (Sequence name “England/CAMB-7526A/2020” dated 12 March 2020) using ape (v2.11.1) (40). Tree visualisation was carried out in R in RStudio using ggtree (v3.8.2) and metadata heatmaps added using gheatmap() and colour scales using RColorBrewer (v1.1-3) (41,42). The phylogenetic analysis was carried out on a QIB cloud server, adapted from CLIMB (29).

### Wave definitions

VOC wave dates were defined as the week where a VOC (Alpha, Delta or Omicron) became dominant (>51.0%) and remained dominant until replaced by another VOC using the full dataset. The waves were named after the dominant VOC. The period of time enclosing lineages before VOC nomenclature began was labelled pre-VOC. For the first and last wave, dates of the beginning and end of the wave respectively correlate to the boundaries of the sequence dataset. Omicron subwaves were also calculated using this definition.

### Statistical modelling

Univariate K-means clustering was used to evaluate temporal clustering of first lineage emergence. Lineages were clustered based the number of days between their first appearance and the reference date (first date of appearance of any lineage; 12 March 2020). The algorithm was implemented in R using Ckmeans.1d.dp (v4.3.5) with the optimal number of clusters estimated based on the Bayesian Information Criterion (BIC) (43).

Network analysis was used to describe the temporal relationship between the first appearances of different lineages. A directed network was constructed with individual lineages represented as nodes; an edge from lineage A to lineage B was established if lineage A was present in the week preceding the first appearance of lineage B. The relationship between longevity and outdegree, where outdegree refers to the number of direct edges originating from a node (indicating the number of lineages that emerged following the presence of a particular lineage in the previous week), was explored by means of linear regression of outdegree on longevity using the R base stats package. Lineages were clustered based on their residuals by means of univariate K-means clustering using the Ckmeans.1d.dp package with a predefined number of clusters, determined through inspection of the residual distribution.

Graphical analysis carried out in ggplot2 and igraph (v2.0.11) (44). The walktrap algorithm was used to identify communities of lineages, implemented through the cluster_walktrap() function in igraph. The optimal number of steps (4) was selected based on the value that maximized modularity. Graphical visualisation was carried out in igraph and ggraph (v2.2.1) (45).

### County comparison analysis

Sequences per county were grouped from the COG-UK metadata using group_by() and count() within dplyr (36). Cumulative COVID-19 case numbers for England by region by specimen date were downloaded from http://coronavirus.data.gov.uk/details/downloads, with ‘Upper Tier Local Authority’ and ‘cumCasesBySpecimenDate’ specified under ‘metrics’ on 16/01/2024. The output was restricted to cases from March 2020 to December 2022, and matched to the number of sequences per county.

Sequencing coverage was calculated as detailed in 2.3, and counties with a 5-10% sequencing coverage were considered for comparison. The median() function within the base stats package was used to calculate median age per county using 2021 census data using a custom dataset with ‘all usual residents’, ‘upper tier local authorities’ and ‘England and Wales’ and ‘Age – 101 categories’ as chosen variables (33). The curated age dataset is available here: https://www.ons.gov.uk/datasets/create/filter-outputs/f79dce51-e02f-44bd-90c9-b78af3f1bd42#get-data. Population density data was downloaded using ‘all usual residents’, ‘Upper tier local authorities’ and ‘England and Wales’ as chosen variables, with the curated population density dataset available here: https://www.ons.gov.uk/datasets/TS006/editions/2021/versions/4/filter-outputs/8a0b9a92-746c-4a7e-9fa8-eed41723356e#get-data. ‘All usual residents’, ‘countries’, ‘England’ were selected to pull the population density for England, with the dataset available here: https://www.ons.gov.uk/datasets/TS006/editions/2021/versions/4/filter-outputs/ecabd869-f0c1-4a02-a5e9-11ebc9fee3e0#get-data. All data from ONS was downloaded on 10/06/2024. Chosen counties Suffolk and Hertfordshire underwent analysis as detailed in ‘Lineage diversity’ and ‘Wave definitions’. A map was created using a UK county shapefile obtained from the Office for National Statistics (https://geoportal.statistics.gov.uk) under Counties and Unitary Authorities (December 2023) (46). This was created in R using packages sf (v1.0-16) and ggplot2 (47).

## Results

### Confirmed cases and sequencing coverage

Between 10 March 2020 and 31 December 2022, there were 296,429 recorded cases of COVID-19 in Norfolk (48). There were a total of 29,406 whole genome SARS-CoV-2 sequences from positive Norfolk-based COVID-19 samples present in the COG-UK dataset, giving a 9.9% sequencing coverage over the study period. The first recorded COVID-19 case was dated 10 March 2020, and the first sequence dated 12 March 2020 (**Figure 1**).

**Figure 1.**
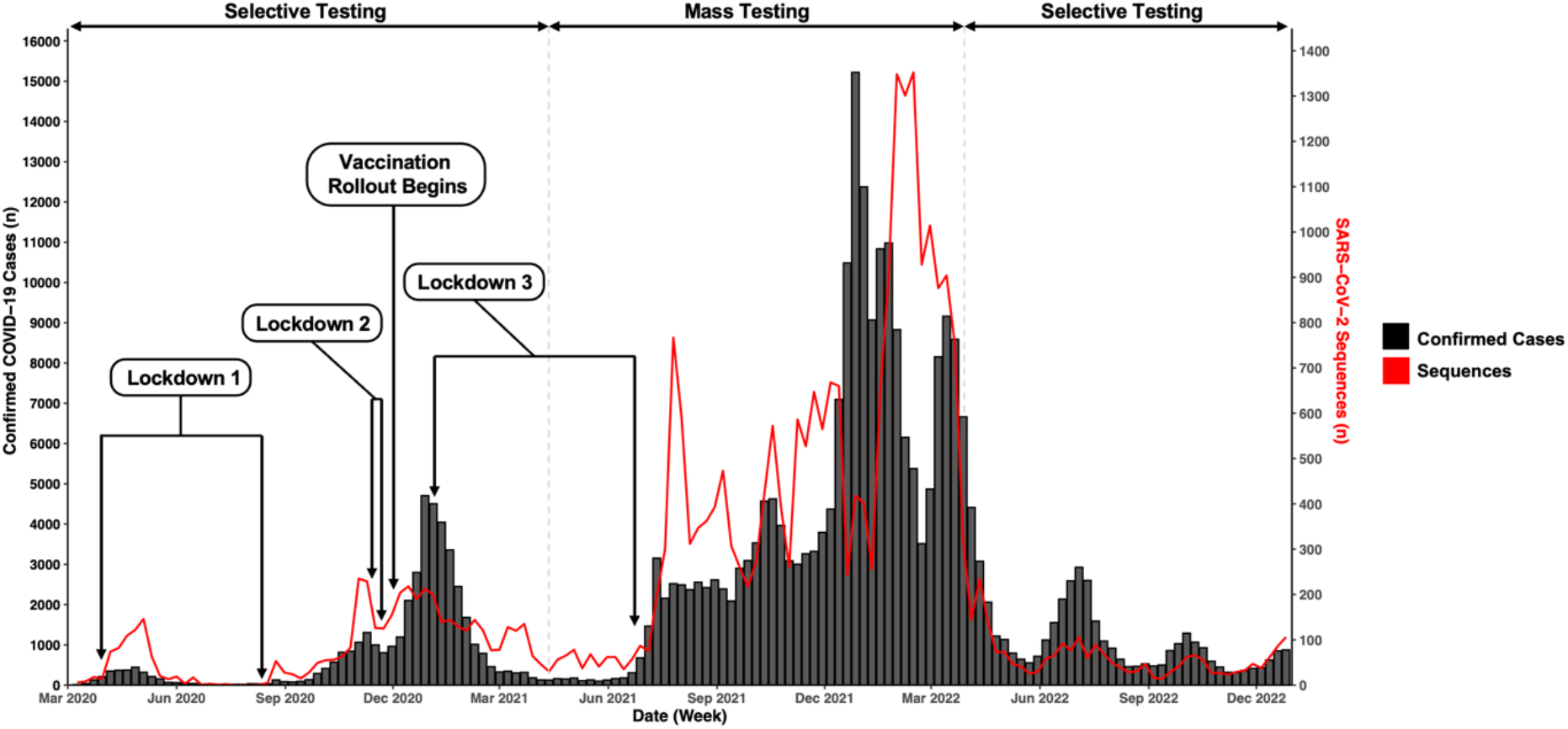
Recorded COVID-19 cases and SARS-CoV-2 sequences in Norfolk, UK, March 2020 – December 2022. Number of recorded positive COVID-19 cases (black, left axis) against the number of SARS-CoV-2 sequences (red, right axis) in Norfolk, UK during the study period. Regional case data obtained from the UK Government COVID-19 dashboard and regional sequence data obtained from COG-UK. Testing data included samples from multiple testing formats including lateral flow devices and polymerase chain reaction tests. Duplicate cases may be present where both were undertaken for the same case, dependent upon national testing requirements. Contextual information regarding lockdowns, vaccines and testing availability was obtained from UK Government press releases (49–55).

### Patient demographics

28,178 (95.8%) of sequences had corresponding patient sex metadata. Of these, 15,369 (54.5%) were from females and 12,788 (45.4%) from males; 21 (0.1%) were labelled ‘other’. 2,974 (10.1%) of sequences had associated patient age metadata, although all age samples had corresponding sex information (**Supplementary Figure 1**). Fewer samples were seen within 0-10 (*n* = 63), 11-20 (*n* = 126) and the 91+ (*n* = 213) age groups. The 81-90 age bracket housed the largest number of samples (*n* = 557).

### Global lineage presence

401 unique global lineages (Pango defined) were sequenced from cases in Norfolk (**Supplementary Table 2**) (3,31). Heterogeneity was seen within the overall number of appearances of each lineage, with each appearing on average 73.3 times (*SD* = 435.1, *SE* = 21.7). 280 (69.8%) appeared more than once and only 125 (31.2%) ten times or more.

The highest number of lineages present in a single week was 37 in the week commencing (w/c) 14 March 2020, followed by 35 in the w/c 2 November 2020 and 13 December 2021(**Figure 3A**). 144 of 147 study weeks contained at least one lineage, with multiple (*n* > 2) co-occurring lineages recorded in 136 weeks within the dataset. Shannon’s Diversity Index (H) was implemented to estimate the diversity per week **(Figure 2A)**, for the most part mirroring the rise and fall of lineage numbers. A consecutive three week period (20 July 2020 - 9 August 2020) was the only period to contain no recorded lineages as no sequences were present for these weeks. Of those with one recorded lineage, a consecutive 5 week period (w/c 15 March 2021 – 18 April 2021) was dominated by lineage B.1.1.7, the Alpha VOC, across 401 samples. The w/c 22 June 2020, 10 August 2020 and 17 August 2020 also had one circulating lineage, but were B.3, B.1.1.347 and B.1.198 respectively – no VOC – and contained 1, 1 and 2 sequences each.

**Figure 2.**
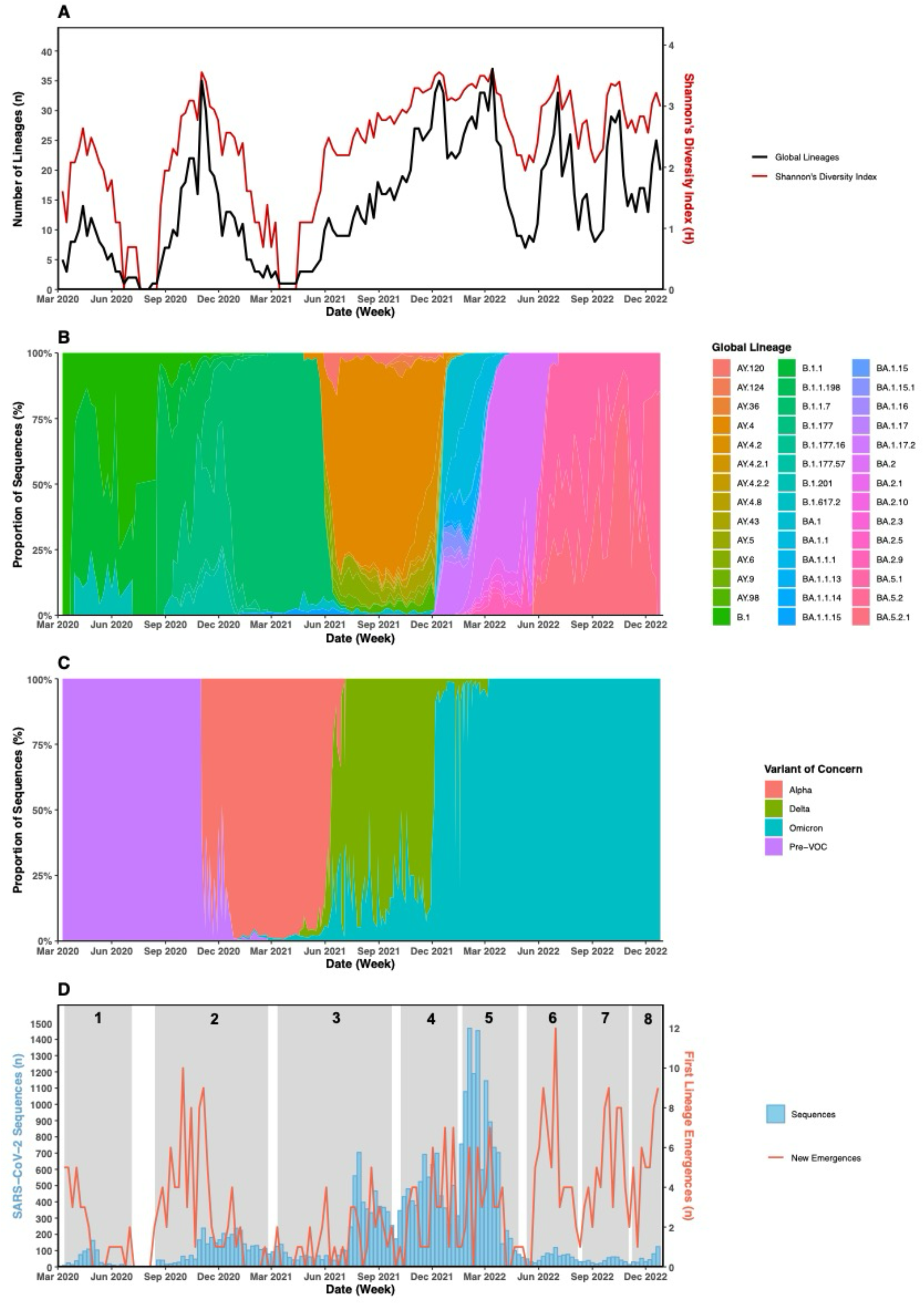
SARS-CoV-2 lineage diversity in Norfolk, UK, March 2020 – December 2022. **(A)** Number of unique SARS-CoV-2 global lineages (black, left axis) co-occurring at weekly intervals in Norfolk throughout the core timeline of the COVID-19 pandemic and the alpha diversity of SARS-CoV-2 each week (red, right axis) using Shannon’s Diversity Index (H). All 401 global lineages represented. **(B)** Proportional appearance of individual SARS-CoV-2 lineages per week across the study period. Focused dataset containing the 42 most commonly appearing lineages shown. Weeks which contain no sequences (30 July 2020 – 09 August 2020) excluded (shown in A) **(C)** Proportional appearance of variants of concern (VOC) per week for the 42 lineage dataset. Those without a VOC designation in the early pandemic fell under the term ‘Pre-VOC’ (purple). Weeks shown in A which contain no sequences excluded (**D)** Univariate k-means temporal clustering of first recorded SARS-CoV-2 lineage emergences. 8 identified chronological periods (clusters) highlighted in grey. Number of first recorded lineage emergences per week (red) against the number of recorded sequences each week (blue).

**Figure 3.**
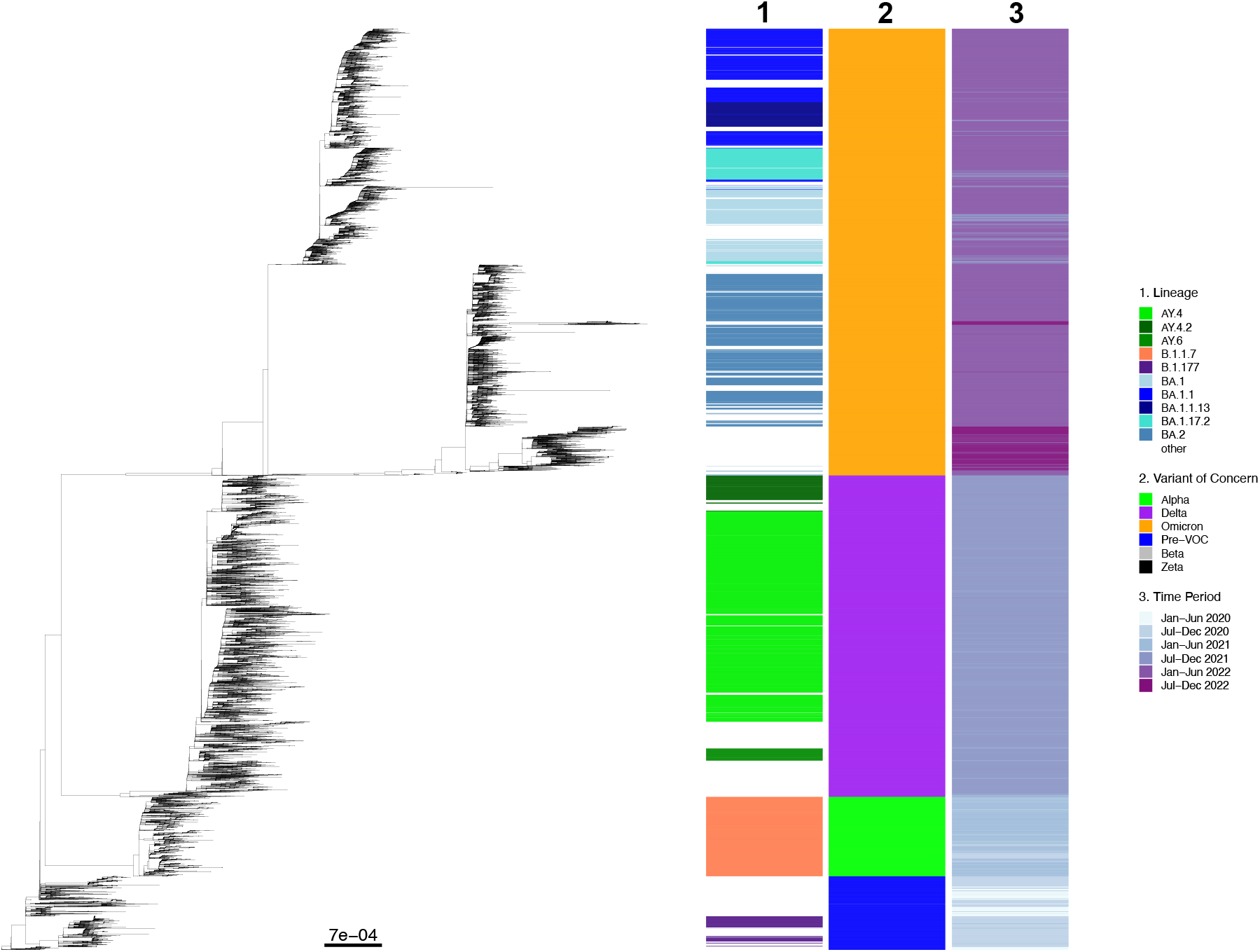
Phylogenetic relationship of SARS-CoV-2 sequences from Norfolk, UK, March 2020 – December 2022. Core SNP ML tree for 29,406 Norfolk-based SARS-CoV-2 sequences downloaded from the public COG-UK dataset, constructed with 7641 formative SNPs. Heatmap 1 represents lineage information for the top ten appearing lineages (n = 20,926 sequences, 71.2% of the overall dataset) with the remaining 319 (n = 8,480 sequences, 28.8% of the overall dataset) were labelled as ‘other’ and not coloured. Heatmap 2 represents variant of concern (VOC) identity for each sequence. Beta and Zeta VOCs were present in low numbers (n= 8 and n=1 respectively) and therefore are not shown. Heatmap 3 represents the time period during which sequencing took place, in 6 month increments from January 2020 to December 2022. Scale is nucleotide substitutions per site.

To assess the most frequently appearing lineages (which had the biggest impact on public health in Norfolk) we focused upon a dataset of 42 lineages (**Table 1**), which accounted for 90.3% (*n* = 26,563) of sequences. The average lineage appearance was 632.5 (*SD* = 1220.8, *SE* = 188.4). The remaining 2,843 sequences belonged to 359 lineages which each appeared on average 7.9 times (*SD* = 11.7, *SE* = 0.6).

**Table 1.** Total lineage appearances within the focused dataset.

| Lineage | Total Appearances (n) | Variant of Concern |
| --- | --- | --- |
| AY.4 | 6540 | Delta |
| BA.2 | 3823 | Omicron |
| BA.1.1 | 2586 | Omicron |
| B.1.1.7 | 2536 | Alpha |
| BA.1 | 1784 | Omicron |
| BA.1.17.2 | 1080 | Omicron |
| AY.4.2 | 849 | Delta |
| BA.1.1.13 | 769 | Omicron |
| B.1.177 | 566 | Pre-VOC |
| AY.6 | 393 | Delta |
| AY.43 | 383 | Delta |
| AY.98 | 341 | Delta |
| BA.2.9 | 318 | Omicron |
| B.1.1 | 308 | Pre-VOC |
| AY.5 | 304 | Delta |
| BA.2.1 | 298 | Omicron |
| BA.1.15.1 | 294 | Omicron |
| B.1.1.198 | 272 | Pre-VOC |
| BA.1.1.14 | 250 | Omicron |
| AY.4.2.2 | 222 | Delta |
| AY.120 | 213 | Delta |
| B.1.177.57 | 199 | Pre-VOC |
| BA.5.1 | 193 | Omicron |
| B.1 | 154 | Pre-VOC |
| BA.1.1.15 | 152 | Omicron |
| BA.2.3 | 141 | Omicron |
| BA.1.17 | 141 | Omicron |
| BA.2.10 | 140 | Omicron |
| BA.5.2.1 | 138 | Omicron |
| BA.1.15 | 133 | Omicron |
| BA.5.2 | 132 | Omicron |
| BA.2.5 | 116 | Omicron |
| B.1.617.2 | 109 | Delta |
| BA.1.16 | 108 | Omicron |
| AY.124 | 93 | Delta |
| AY.4.2.1 | 79 | Delta |
| AY.4.8 | 73 | Delta |
| AY.9 | 70 | Delta |
| BA.1.1.1 | 69 | Omicron |
| AY.36 | 68 | Delta |
| B.1.201 | 63 | Pre-VOC |
| B.1.177.16 | 63 | Pre-VOC |

*SARS-CoV-2 global lineages observed within Norfolk, UK March 2020 – December 2022 showing the 42 most commonly appearing lineages in descending order of appearance. 'Pre-VOC' refers to lineages appearing in the early pandemic before variant of concern (VOC) naming conventions began in May 2021. The number of appearances can also be defined as the number of cases per lineage.*

### Variant of concern presence

93.9% (*n =* 24,938) of sequences in the focused dataset had a VOC designation (**Table 2**). All sequences which did not have a VOC designation (*n* = 1625) appeared in the early pandemic before VOC designation began (May 2021 onwards) and were thus labelled ‘pre-VOC’. Alpha, Delta and Omicron VOCs were present, with Omicron accounting for almost half (12,665 [47.7%]) of the sequences. Delta and Alpha accounted for 9,737 (36.7%) and 2,536 (9.6%) sequences respectively.

**Table 2.** Global variant of concern presence in Norfolk, UK between March 2020 and December 2022.

| Variant of Concern | Number of Lineages (n) | Number of Sequences (n) | Percentage of Total Sequences (%) |
| --- | --- | --- | --- |
| Pre-VOC | 7 | 1625 | 6.1 |
| Alpha | 1 | 2536 | 9.6 |
| Delta | 14 | 9737 | 36.7 |
| Omicron | 20 | 12665 | 47.7 |

*Number of unique SARS-CoV-2 lineages and sequences under each variant of concern (VOC) designation observed in 42 lineage dataset.*

The number of unique lineages identified in each VOC bracket increased post-VOC designation (**Table 1**). Pre-VOC designation, 7 lineages were circulating. Alpha only contained one unique lineage (B.1.1.7), whereas Delta and Omicron contained 14 and 20 respectively, reflecting an increase in SARS-CoV-2 diversity with pandemic progression.

### Lineage diversity

SARS-CoV-2 diversified throughout the pandemic, with lineages transiently displaying patterns of emergence, persistence and then extinction when a lineage was outcompeted by a succeeding lineage (**Figure 2B**). We constructed a core single nucleotide polymorphism (SNP) maximum likelihood (ML) tree to display the phylogenetic relationship between all sequences during the study period (**Figure 3**).The number of days a lineage was circulating (the longevity) varied, with no lineage reappearance observed after extinction. In the full dataset containing 401 lineages, lineages were present over an average of 51.1 days (*SD =* 62.9, *SE =* 3.1) with sequences recorded on average 13.3 (*SD =* 27.7, *SE =* 1.4) days within this span. The minimum number of days a lineage was present for was 1 day, and the maximum 447 showing considerable lineage-dependent variation. For the dominant 42 lineages, the average longevity was 164.4 days (*SD =* 80.2, *SE =* 12.4) with recorded sequences on an average of 80.9 days (*SD =* 42.0, *SE =* 6.5). However, the minimum longevity within this dataset was 58 days, highlighting lineages with a greater impact on public health were circulating for longer time periods.

### Variant of concern diversity and epidemic waves

Four VOC epidemic waves were present across the full dataset (**Figure 2C**), with dated boundaries (**Table 4**) defined by the week the overall presence of one or more lineages falling under the respective VOC designation totalled and remained >51.0% until outcompeted by a succeeding VOC. The VOCs displayed the same transient nature as the lineage level. These waves began with lineages with pre-VOC designation lineages in the early pandemic, followed by Alpha, Delta and Omicron VOCs.

**Table 4.** Variant of concern wave dates, length and global lineage presence.

| VOC Wave | Date Range (week commencing) | Length (week) | Number of sequences (n) | Total Global Lineages (n) | Most Frequently Appearing Lineage | % of Sequences Within VOC wave (%) |
| --- | --- | --- | --- | --- | --- | --- |
| Pre-VOC | 09-Mar-20* to 21-Dec-20 | 42 | 1946 | 8 | B.1.177 | 21.6 |
| Alpha | 28-Dec-20 to 24-May-21 | 22 | 2204 | 11 | B.1.1.7 | 95.6 |
| Delta | 31-May-21 to 13-Dec-21 | 29 | 9890 | 23 | AY.4 | 65.5 |
| Omicron | 20-Dec-21 to 26-Dec-22* | 54 | 12563 | 31 | BA.2 | 30.5 |

*Dated range and length of epidemic VOC waves and the number of unique SARS-CoV-2 lineages present within the bounds of each wave. 20 = 2020, 21 = 2021, 22 = 2022. \* = start/end of study period*

The crossover period between each wave, defined as where an emerging VOC rose in dominance and superseded the dominating VOC, varied (**Table 4**). The Alpha variant took 8 weeks from emergence (B.1.1.7, 2 November 2021) in the region to become the dominant VOC, during which B.1.177 (and related lineages - Pre-VOC) and B.1.1.7 (Alpha) lineages were most frequently observed, with B.1.1.7 slowly overtaking in dominance before establishing the Alpha wave in late December. The Delta variant took 6 weeks to establish itself as the dominant VOC, after emergence in the region on 20 April 2021, during which B.1.1.7 slowly decreased, superseded by multiple Delta lineages. The Omicron variant briefly appeared in the region on the 3 January 2021 (BA.1.1.15) and once again 13 January 2021 (BA.1.1.529) with the next appearance on 4 December 2021, 11 months later. It became dominant on the 20^th^ December 2021, 3 weeks after the last emergence of the variant in the region, and represented the most rapid crossover period of all VOCs.

The total number of global lineages increased with each consecutive wave (**Table 4, Supplementary Tables 3-6)**. Each wave contained lineages from the dominating VOC, alongside those from neighbouring VOC waves, accounting for crossover periods. After the pre-VOC wave, the length of each VOC wave increased with the Alpha wave lasting 22 weeks and the Delta wave 29 weeks. Omicron then dominated for at least 54 weeks (**Table 4**), beginning in the w/c 20 December 2021 until the end of the study period.

In the pre-VOC wave, there was no single consistent predominant lineage (**Figure 2B**), however

B.1.177 appeared most frequently during this time period, closely followed by B.1.1.7 (the Alpha variant) which appeared in high numbers due to an 8 week crossover period seen between the two epidemic waves (**Table 4**). The Alpha and Delta waves were dominated by B.1.1.7 and AY.4 respectively. The Omicron wave did not have a single dominant lineage, but rather three that were proportionately the most frequent during the wave; BA.2, BA.1.1 and BA.1 were present in 30.5%, 20.5% and 13.3% of sequences in this wave respectively. Of the 20 Omicron lineages present in this wave, only 5 were present in more than 5.0% of samples, displaying higher variability within the Omicron VOC compared to previous VOCs (**Supplementary Table 6**).

Due to the significant length of the Omicron epidemic wave (54 weeks), we sought to identify subwaves within this period derived from larger parental sublineages. Three waves were identified from sublineages BA.1, BA.2 and BA.5. These subwaves spanned 44 of 54 weeks of the Omicron epidemic wave (**Table 5**), with the remaining weeks at the end of the wave where there was no consistently dominating sublineage. BA.5, BQ.1 and XBB parental lineages were present during this time but not in large enough numbers to sustain a subwave.

**Table 5.** Subwaves present within the Omicron epidemic wave.

| Subwave parental lineage | Date Range<br>(week commencing) | Length<br>(week) |
| --- | --- | --- |
| BA.1 | 20-Dec-21 to 21-Feb-22 | 11 |
| BA.2 | 28-Feb-22 to 30-May-22 | 13 |
| BA.5 | 06-Jun-22 to 17-Oct-22 | 20 |
| Undefined | 24-Oct-22 to 26-Dec-22* | 10 |

*Dated range and length of subwaves within the Omicron VOC epidemic wave. The end date for the Omicron wave refers to the boundary collection dates of the sequencing data, and therefore does not reflect the true end date of the Omicron epidemic wave. Undefined refers to a 10 week period at the end of the wave where no sublineage group was present in >51% of sequences. 21 = 2021, 22 = 2022. \* = end of study period*

Omicron displayed different evolutionary dynamics (**Figure 3**) to other VOCs. Three distinct clades were present, with lineages from parental groups BA.1, BA.2 and a third containing BA.4, BA.5 and BQ.1 (sublineage of BA.5). The BA.2 clade contained a longer branch containing XBB sublineages (a BA.2 hybrid lineage), displaying differing evolutionary patterns within each distinct clade.

### Statistical modelling

We applied statistical modelling to the dataset to see if trends complementary to using genomic epidemiology methodology could be established, relevant to pandemic response. To find periods of time when emergence of new lineages clustered, we applied univariate k-means temporal clustering to the time to emergence in the full dataset containing all 401 lineages. 8 clusters were identified chronologically spanning the length of the pandemic (**Figure 2D)**, each comprising on average 50.1 lineages (*SD =* 17.6, *SE =* 6.0) that correspond to sequential VOC presence, accounting for the delay between first lineage emergence and an increase in cases caused by the respective lineage (**Figure 2B, Supplementary Table 2)**. In particular, clusters 5-8 contain the first lineage emergences of major Omicron sublineages (BA.1, BA.2, BA.4, BA.5, BQ.1 and XBB) which contribute to subwaves previously detailed.

We assessed the relationship between lineage longevity and first recorded lineage emergence using a directional network analysis. We defined lineages as nodes and edges were defined as existing between lineage A and B when lineage A was present the week before the emergence of lineage B. 19,433 edges originating from 215 nodes were identified within the dataset (**Supplementary Figure 2A**). The average number of edges originating from a singular node (the outdegree) was 90.4 (*SD =* 75.0, *SE =* 5.1), displaying high yet variable correlation between lineage longevity and outdegree. 81 lineages had an outdegree over 100.

A linear regression between longevity and the number of outdegrees indicated, as expected, a strong correlation between longevity and outdegree (r=0.64). However, visual exploration showed that the relationship was heterogeneous with 3 clusters (**Supplementary Figure 3**) of differing correlation strengths were identified within the dataset, with differing number of lineages present in each (*n =* 25, *n =* 286, *n =* 90). Cluster 1 included lineages for which there was no clear relationship between longevity and the number of outdegree, Cluster 2 a mild correlation and Cluster 3 a strong correlation. 8 of the top 10 appearing lineages (**Table 1**) were present in Cluster 1, aside from AY.6 and BA.2, which appeared in Cluster 3 and Cluster 2 respectively. The relationship here may be explained by a long overall longevity but fewer appearances within this or limited emergences during their longevity, reducing the outdegree.

To explore whether impactful groups of lineages (i.e. VOC groups) could be identified within the network, a walktrap algorithm was applied to find connected subgroups within the network which housed stronger relationships. Six nested subgroups (named communities, **Supplementary Table 2, Supplementary Figure 3B-G**) were identified with all lineages represented. All identified communities contained Pre-VOC lineages alongside at least one VOC group. Community 5 contained all major VOCs observed in the region – Alpha, Delta, Omicron – alongside Pre-VOC lineages.

Community 2 and 6 contained Pre-VOC and Omicron lineages, with the remaining communities (1, 3 and 4) containing Pre-VOC, Omicron and Delta lineages. Major sublineages within each VOC group were distributed across multiple communities, with no clear descriptive correlation.

### Regional comparison

To determine whether SARS-CoV-2 lineages and VOC presence in Norfolk were reflective of the national picture during the COVID-19 pandemic we compared our findings to other geographical regions in the UK (**Supplementary Figure 4**). Given Norfolk is largely coastal, rural and has a relatively low population density (170.2 people per km^2^ in comparison to England’s population density of 433.5 people per km^2^), we sought to see if these county demographics had an impact on genomic epidemiology. We selected one neighbouring region with similar demographics (Suffolk, 200.2 people per km^2^) and one non-bordering county with a higher population density and close proximity links to London (Hertfordshire, 729.6 people per km^2^). Median age of the population was also considered, with Norfolk having a slightly older population of 45 years, Suffolk 45 years and Hertfordshire 40 years, comparable to the UK median (40 years). These regions also had similar sequencing coverage to Norfolk (9.9%) within the COG-UK dataset (Suffolk = 9.7%, Hertfordshire = 10.7%).

Similar patterns of transient lineage emergence, persistence and extinction were present across the three regions at both the lineage and VOC level, however the temporal presence of individual lineages varied in comparison to Norfolk (**Figure 4**). The dominant lineage within each VOC wave was the same in all regions for Alpha (B.1.1.7), Delta (AY.4) and Omicron (BA.2) **(Supplementary Table 7)**. In the Pre-VOC wave, lineage B.1.177 was the predominant lineage in both Norfolk and Suffolk, and B.1.1 predominant in Hertfordshire.

**Figure 4.**
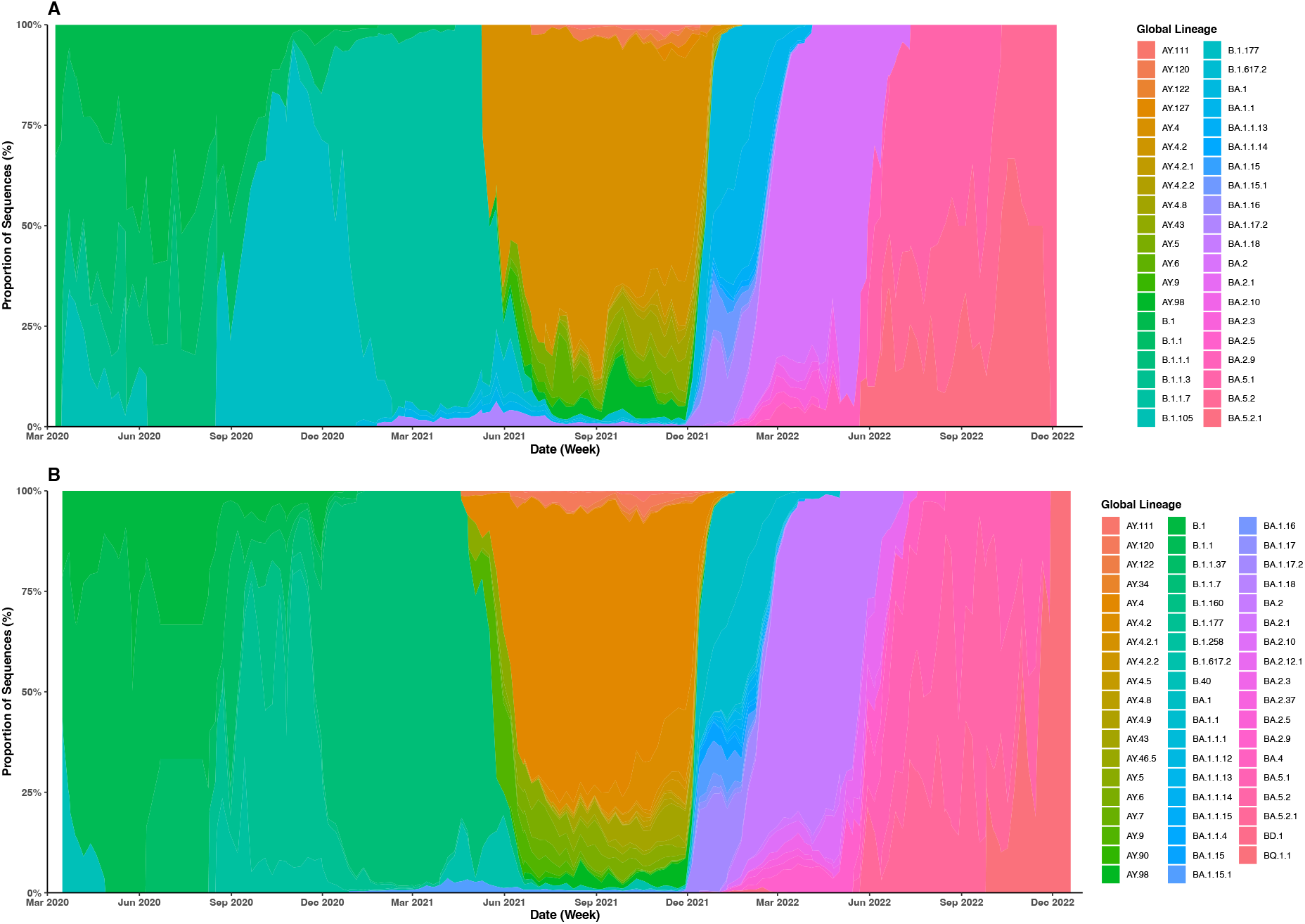
Lineage diversity per week in comparison counties, March 2020 – December 2022 (A) Proportional appearance of individual SARS-CoV-2 lineages across the COVID-19 pandemic in Suffolk, UK. **(B)** Proportional appearance of individual SARS-CoV-2 lineages across the COVID-19 pandemic in Hertfordshire, UK. Those without a VOC designation in the early pandemic fell under the term ‘Pre-VOC’ (purple). Focused datasets from both regions focused to show the most commonly appearing lineages.

Across the regions, 721 unique global lineages were present, with 236 common to all. Norfolk and Suffolk shared a similar level of viral diversity, with 399 global lineages present in Suffolk compared to 401 in Norfolk. Hertfordshire had more diversity, with 554 recorded global lineages present.

Hertfordshire had the most unique lineages, followed by Norfolk and Suffolk (*n* = 202, *n* = 70, *n* = 53). Of the common lineages, Norfolk shared similar levels with both Suffolk and Hertfordshire (*n* = 280, *n* = 286), however a greater overlap was seen between Suffolk and Hertfordshire (*n* = 301).

Norfolk and Hertfordshire shared the most unique lineages which were common to the two counties but not Suffolk (*n* = 121), followed by Suffolk and Hertfordshire (*n* = 65) and finally Norfolk and Suffolk housed the fewest lineages unique to the two counties (*n =* 44).

In comparison to Norfolk, all waves in Hertfordshire started and ended earlier (**Table 6, Supplementary Figure 5**). Suffolk dates were more variable, with the Alpha and Omicron waves starting later, and Delta starting earlier. In terms of wavelength, the Pre-VOC wave was a month shorter in Hertfordshire, with Alpha taking over earlier and the wave lasting two weeks longer. The Pre-VOC wave was also a week longer in Suffolk, and the Delta wave three weeks longer.

**Table 6.** VOC wave dates for Norfolk and comparison counties.

|  | Pre-VOC |  | Alpha |  | Delta |  | Omicron |  |
| --- | --- | --- | --- | --- | --- | --- | --- | --- |
|  | Start Date (w/c) | End Date (w/c) | Start Date (w/c) | End Date (w/c) | Start Date (w/c) | End Date (w/c) | Start Date (w/c) | End Date (w/c) |
| Norfolk | 09-Mar-20* | 21-Dec-20 | 28-Dec-20 | 24-May-21 | 31-May-21 | 13-Dec-21 | 20-Dec-21 | 26-Dec-22* |
| Suffolk | 09-Mar-20* | 28-Dec-20 | 04-Jan-21 | 10-May-21 | 17-May-21 | 20-Dec-21 | 27-Dec-21 | 26-Dec-22* |
| Hertfordshire | 16-Mar-20* | 30-Nov-20 | 07-Dec-20 | 17-May-21 | 24-May-21 | 06-Dec-21 | 13-Dec-21 | 26-Dec-22* |

Dated range and length of waves for Norfolk and comparison counties Suffolk and Hertfordshire. 20 = 2020, 21 = 2021, 22 = 2022. \* = start/end of study data.

## Discussion

We have examined the genomic epidemiology of SARS-CoV-2 in one densely sequenced UK region over the core timeframe of the pandemic, giving retrospective insight of a pandemic virus at a local scale. SARS-CoV-2 was diverse in Norfolk, with 401 global lineages present across the study period. Despite this, 10% of lineages accounted for 90.3% of the total number of sequences, showing a highly skewed heterogeneous population. These lineages formed our focused dataset, consisting of 42 dominant lineages, focusing analysis on the lineages which were the most prevalent and had the most significant impact on regional public health. However, this does not indicate that the remaining 90% of lineages were insignificant; these give insight into genetic diversity and may include fitness enhancing mutations which may have contributed to a subsequent dominant lineage. These may also represent small, isolated regional or inter-regional transmission clusters.

Our application of statistical modelling to the dataset, looking specifically at first lineage emergences, identified 8 chronological periods of temporal clusters of lineage emergence, which roughly correlated to identified VOC waves and significant sublineages within these. This suggests that this methodology may be complementary to genomic identification of significant time periods within a pandemic setting. However, our further analysis using a network in which lineages were simply linked if the emergence of one lineage was preceded by the presence of another in the previous week, did not correlate to genomic results with identified communities not representing phylogenetically relevant groups. We used first lineage emergence as a metric, which should be noted as the first detected emergence of a lineage in the region, therefore it is possible that lineages were circulating before their first detection regionally. This may explain why groupings differ from phylogenetic relatedness, as first detection does not reflect stepwise viral mutation. Other metrics may be more suitable and require further investigation.

As a county, Norfolk is a largely rural, coastal region in the East of England, bordered by two neighbouring counties: Suffolk and Cambridgeshire, forming East Anglia. The mixture of rural and urban areas are reflective of the wider demographics of the UK, however with a relatively low regional population density. During the study period (reported in the 2021 UK census), Norfolk had a population of 916,120, composed of 51.0% females and 49.0% males (56). The county has a higher percentage of those aged over 65, with 24.4% compared to 18.4% for England overall (56,57). Inter-regional variation was seen between SARS-CoV-2 lineages in both neighbouring (Norfolk and Suffolk) and more distant (Norfolk and Hertfordshire) counties. Norfolk’s VOC waves were consistently behind Hertfordshire, highlighting a delay in viral dissemination from more distant, densely populated areas, likely dependent on population travel patterns. The increased lineage diversity seen in Hertfordshire in comparison to both Norfolk and Suffolk may be explained by regional proximity to Greater London, giving rise to potential introduction events through commuting, and further possibilities of transmission with domestic and international tourism-related clusters.

Despite the variability seen in both lineage diversity and VOC wave dates between Norfolk and Suffolk, the most commonly appearing lineages within each wave remained the same, with the same patterns of heterogeneity observed, suggesting populational factors causing differing transmission clusters between counties may play a role in establishment of a dominant VOC.

Our findings show a substantial increase in SARS-CoV-2 diversity in Norfolk since the last published report, where 26 global lineages were identified between March – August 2020 (28). In comparison, there were 42 lineages circulating in the COG-UK dataset during this period with only 8 lineages present across both datasets. Our data confirms that during this period B.1.1 was the most common lineage. We identified lineages of AD, B and C designation in COG-UK dataset for this 6 month period, whereas only lineages of B designation were previously observed. Differences may be accounted for through differing sampling intent (i.e. outbreak analysis), sensitivity of data used or updated nomenclature since publication.

Further studies across East Anglia have reported lineage circulation, with presence of those in Norfolk at the same time indicating inter-county transmission. B.1.60.7, B.1.177 and B.1.177.16 were identified in Cambridge between October – December 2022 and were present in Norfolk at the same time (58). In February – May 2020, B.1, B.1.1, B.1.1.1 B.1.10 and B.1.1.3 were found to be circulating both in East Anglian care facilities and in Norfolk (59). B, B.1, B.1.1, B.1.1.10 and B.3 were found to be circulating in a further set of East Anglian care facilities between March – April 2020, and were also observed in our dataset (60). However, not all lineages identified in these studies were observed in Norfolk, suggesting differing transmission clusters were present in each location.

VOC epidemic wave dates for COVID-19 infection nationally or regionally by county for the UK have not been established. However, reactive wave dates determined in near real time during the pandemic were defined by a sustained increase in transmission and infection based on metrics including the reproduction rate, growth rate and positivity rate of COVID-19 (61). Our definition, focusing on sequencing data, of VOC epidemic waves indicated the periods during which SARS-CoV-2 variants with elevated risk to public health were dominantly circulating. This reflects SARS-CoV-2 diversity more widely, as these dates consider changes to the virus rather than estimated infection levels. ‘Variant periods’ have been reported by the Office for National Statistics for the UK, split by dates where a VOC was present in over 60% of sequences (12). Despite our use of >51.0% to determine dated wave boundaries, VOC wave dates for Norfolk remain the same when adjusted to >60.0%.

The Alpha and Delta VOC epidemic waves in Norfolk were two weeks behind these national variant period estimates, but Omicron epidemic wave dates were matched; however differences were seen in subwaves (12). BA.1, BA.2 and BA.5 subwaves overlapped with these estimates, with the BA.2 subwave ending a week earlier. Norfolk’s BA.5 subwave lasted 20 weeks and ended in the w/c 17 October 2022, however after this period BA.5, BQ.1 and XBB sublineages were circulating but there was no dominant regional sublineage. BQ.1 and XBB sublineages were reported to be predominantly present in UK cases by January 2023, indicating this time is likely to have been a crossover period between dominant Omicron sublineages (62). Differences between regional Norfolk and national patterns indicate that, once averaged at the national level, resolution from local variation is lost.

The first official recording of the Omicron variant in the UK was on the 27^th^ of November 2021 (17). We noted two appearances of Omicron lineages before this date – BA.1.1.15 on 3 January 2021 and B.1.1.159 on 13 January 2021. The next record after this was 4 December 2021, 11 months later. This is likely due to similar mutational patterns in these sequences which were not designated as Omicron until much later, reclassification of lineages or isolated appearances due to lockdown restrictions (i.e. a reduced opportunity to spread).

The number of sequences present within the COG-UK dataset was impacted by availability of NHS polymerase chain reaction (PCR)-based COVID-19 testing, which changed throughout the study period. UK testing consisted of either PCR test in isolation or an initial positive on a lateral flow device (LFT) followed by a confirmatory PCR test, with positive PCR results forwarded for sequencing. After January 2022, a PCR test was not required after a LFT test, reducing the number of samples for sequencing (49). In the early pandemic, testing was only available to those in clinical care settings, healthcare staff and essential workers (63,64). Mass testing was available between April 2021 and April 2022 but after this period, testing reverted to those with clinical need and healthcare workers (50,51). National lockdowns also impacted sequence numbers, with a consecutive three week period (20 July 2020 – 9 August 2020) with no recorded cases, alongside significantly fewer cases during the first and third lockdowns (**Figure 1**). Despite this fluctuation, substantial SARS-CoV-2 diversity was still captured.

Our study details the genomic epidemiology of SARS-CoV-2 in Norfolk from March 2020 to December 2022, observing significant impacts from few lineages within a diverse population, with their VOC identity forming four epidemic waves. Our findings highlight the important role of incorporating genomic methods in pandemic response and their ability to provide detailed regional insights.

## Supporting information

COG-UK Consortium Author List

## Ethical approval and consent to participate

The use of COG-UK data within this study falls under ethical approval to COG-UK (PHE R&D Ref: NR0195). Further ethical approval was awarded for this study by the University of East Anglia Faculty of Medicine and Health Sciences Research Ethics Subcommittee under reference ETH2223-2684.

## Author contributions

EHH contributed to study design and analysis, and wrote the original draft of the manuscript. JG designed and completed the statistical modelling analysis and contributed to writing the corresponding sections in the manuscript. AJP conceptualised the study and facilitated access to data. GCL, AJP and RAK supervised the project and contributed to study design. All authors contributed to reviewing the manuscript, and GCL edited the manuscript. All authors approved the final version of the manuscript.

## Acknowledgements

We would like to thank COG-UK for their action during the pandemic, and for curating the dataset accessed for this study. A full list of contributors to the COG-UK consortium is available in the supplementary material. We would also like to thank the Core Bioinformatics team at the Quadram Institute, particularly Dr Leonardo de Oliveira Martins, for both technical and phylogenetic analysis support. We would also like to thank Miss Alice Nisbet for providing baseline code for phylogenetic visualisation.

