## Supplementary Figures 1-4, Supplementary Tables 2-7 for "Genomic Epidemiology of SARS-CoV-2 in Norfolk, UK, March 2020 – December 2022"

### Supplementary Material

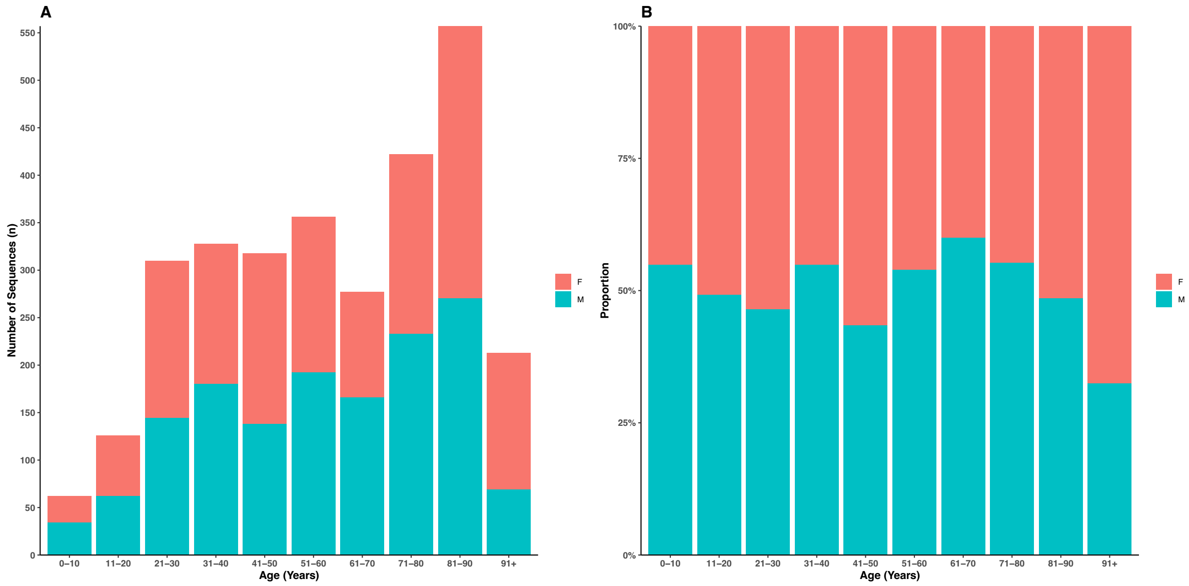


**Supplementary Figure 1. Age and sex distribution of patients where metadata was present for both variables**

*Age and sex metadata from 2,974 patients associated with COG-UK sequencing metadata regionally in Norfolk.* ***A -*** *count of each age band, with* ***B*** *displaying data proportionately. 'Other' group not shown.*

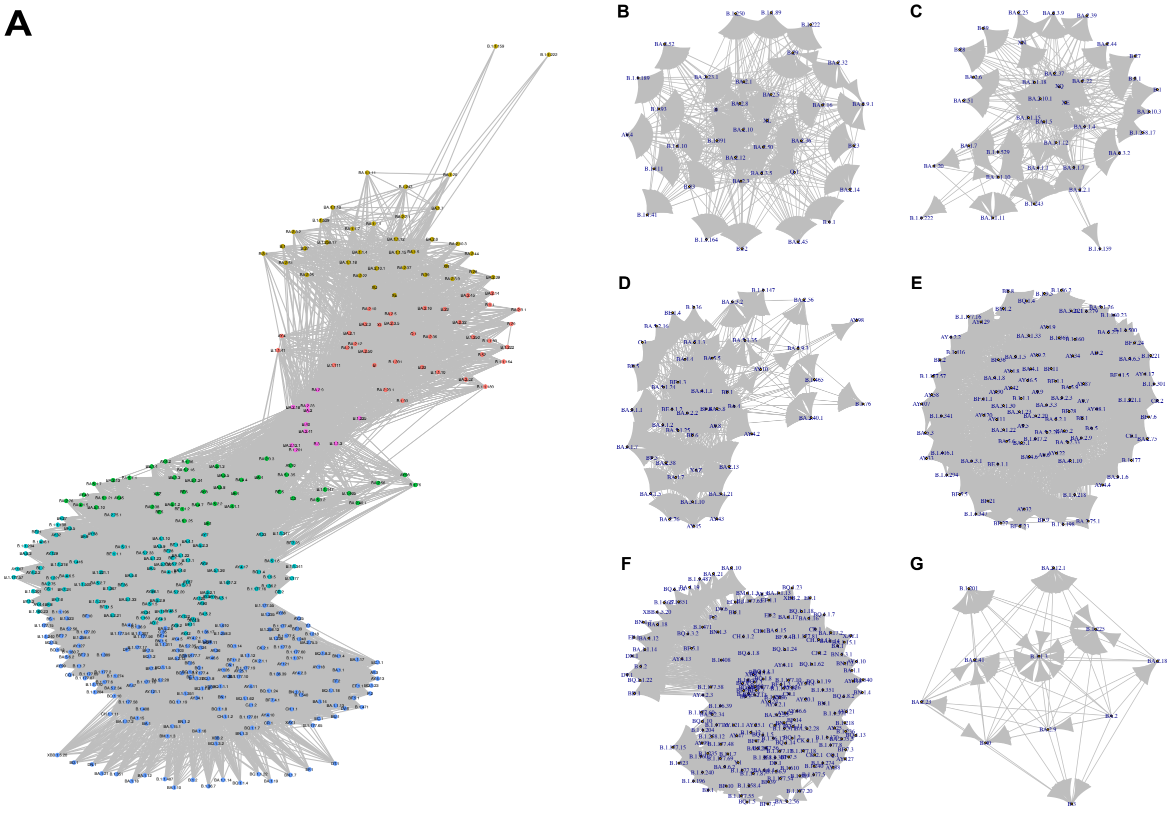


**Supplementary Figure 2. Lineage networks based off first lineage emergences**
***(A)*** *Overall network displaying connectivity between lineages where A appeared the week before B.* ***(B-G)*** *Individual identified subgroups (communities) from within A which housed strong links.*

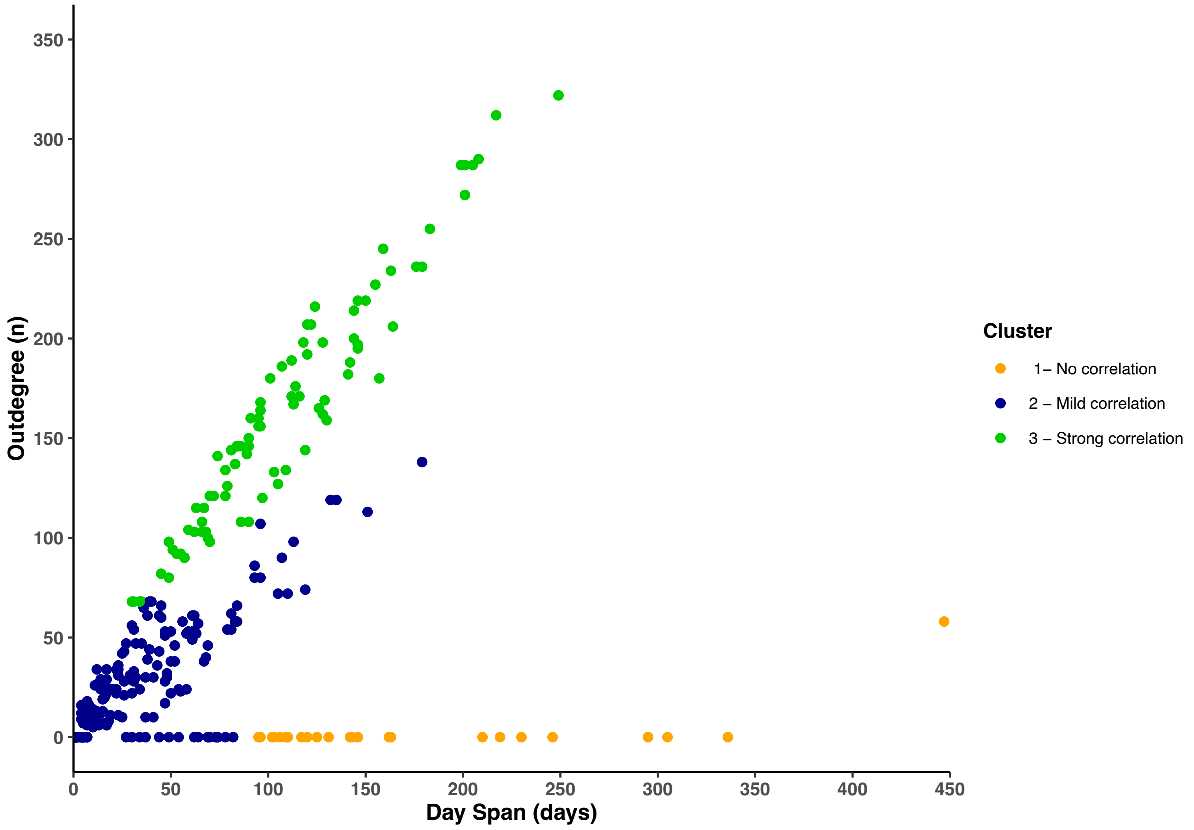


**Supplementary Figure 3. Relationship between longevity and first lineage emergence**

*The total longevity of each lineage plotted against the total number of edges present for each lineage (the outdegree) in the network analysis. Applied K-means clustering identified three clusters of varying correlation strength (no, mild, strong)*

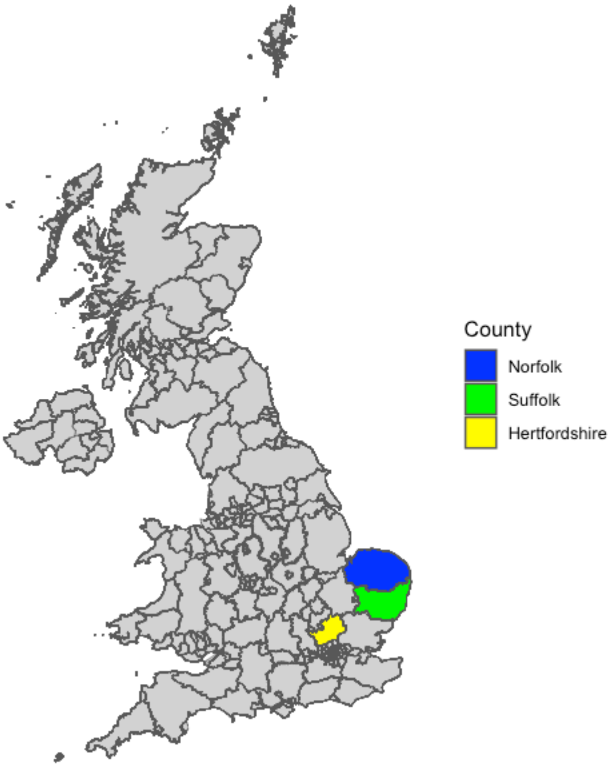


**Supplementary Figure 4. Selected counties for genomic epidemiology comparison metrics**
*Map of the United Kingdom colourised with Norfolk (primary study location, blue) and comparison counties Suffolk (green) and Hertfordshire (yellow).*

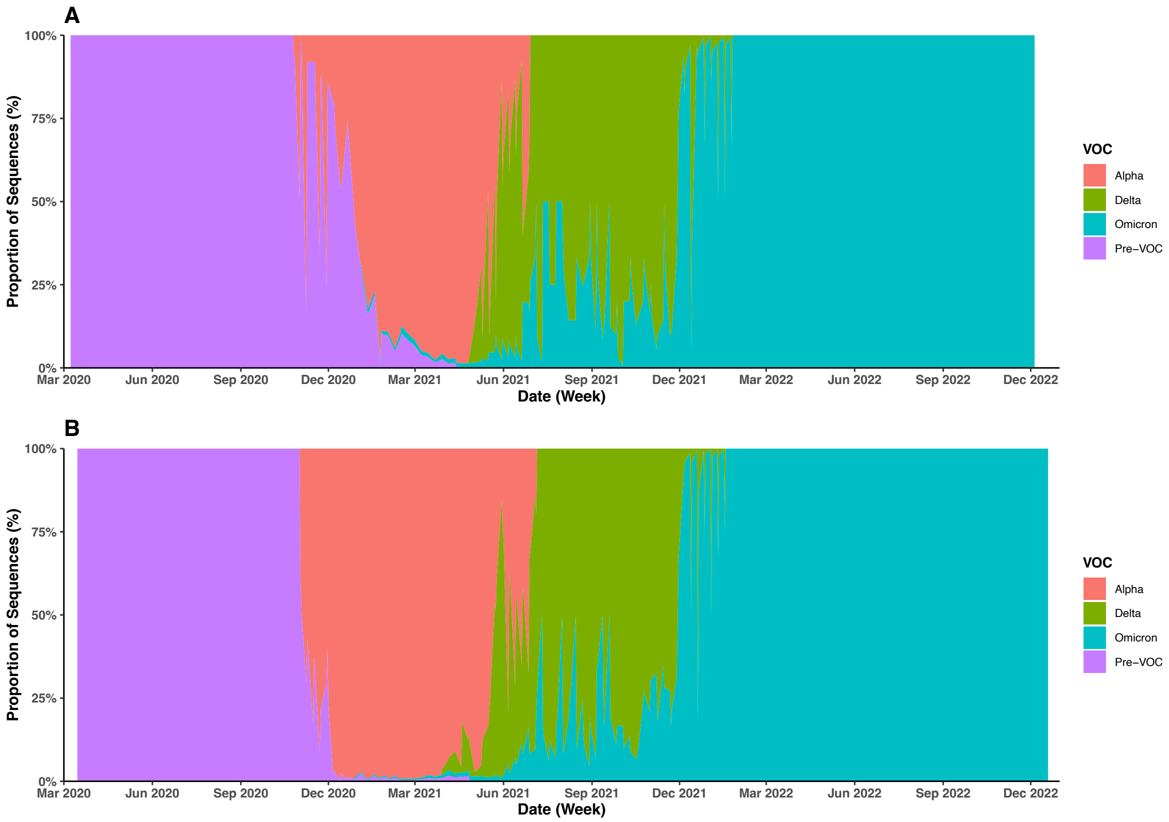


**Supplementary Figure 5. VOC graphs for Suffolk and Hertfordshire**
*Proportional appearance of individual SARS-CoV-2 lineages across the COVID-19 pandemic in (**A**) – Suffolk and (**B**) - Hertfordshire. Lineages in the early pandemic without a VOC designation fall under the term 'Pre-VOC' (purple). Datasets from both regions focused to show the most commonly appearing lineages (top 10th percentile of overall number of appearances).*

**Supplementary Table 2. Total lineage appearances and VOC designation within the full dataset**

| Lineage | Total Appearances (n) | VOC Designation | First Lineage Emergence Temporal Cluster | Longevity/Outdegree Cluster | Assigned Walktrap Community |
|---|---|---|---|---|---|
| AY.4 | 6540 | Delta | 3 | 1 | 2 |
| BA.2 | 3823 | Omicron | 4 | 2 | 2 |
| BA.1.1 | 2586 | Omicron | 4 | 1 | 5 |
| B.1.1.7 | 2536 | Alpha | 2 | 1 | 5 |
| BA.1 | 1784 | Omicron | 4 | 1 | 5 |
| BA.1.17.2 | 1080 | Omicron | 4 | 1 | 5 |
| AY.4.2 | 849 | Delta | 3 | 1 | 2 |
| BA.1.1.13 | 769 | Omicron | 4 | 1 | 5 |
| B.1.177 | 566 | Pre-VOC | 2 | 1 | 5 |
| AY.6 | 393 | Delta | 3 | 3 | 4 |
| AY.43 | 383 | Delta | 3 | 1 | 4 |
| AY.98 | 341 | Delta | 3 | 1 | 5 |
| BA.2.9 | 318 | Omicron | 5 | 2 | 3 |
| B.1.1 | 308 | Pre-VOC | 1 | 1 | 4 |
| AY.5 | 304 | Delta | 3 | 3 | 5 |
| BA.2.1 | 298 | Omicron | 5 | 2 | 6 |
| BA.1.15.1 | 294 | Omicron | 4 | 1 | 5 |
| B.1.1.198 | 272 | Pre-VOC | 2 | 1 | 5 |
| BA.1.1.14 | 250 | Omicron | 4 | 1 | 2 |
| AY.4.2.2 | 222 | Delta | 3 | 1 | 1 |
| AY.120 | 213 | Delta | 3 | 3 | 1 |
| B.1.177.57 | 199 | Pre-VOC | 2 | 1 | 5 |
| BA.5.1 | 193 | Omicron | 6 | 3 | 3 |
| B.1 | 154 | Pre-VOC | 1 | 1 | 4 |
| BA.1.1.15 | 152 | Omicron | 2 | 1 | 2 |
| BA.1.17 | 141 | Omicron | 4 | 1 | 5 |
| BA.2.3 | 141 | Omicron | 5 | 2 | 2 |
| BA.2.10 | 140 | Omicron | 4 | 2 | 1 |
| BA.5.2.1 | 138 | Omicron | 6 | 3 | 4 |
| BA.1.15 | 133 | Omicron | 4 | 1 | 5 |
| BA.5.2 | 132 | Omicron | 6 | 3 | 3 |
| BA.2.5 | 116 | Omicron | 5 | 2 | 6 |
| B.1.617.2 | 109 | Delta | 3 | 3 | 1 |
| BA.1.16 | 108 | Omicron | 4 | 2 | 5 |
| AY.124 | 93 | Delta | 3 | 3 | 1 |
| AY.4.2.1 | 79 | Delta | 3 | 3 | 6 |
| AY.4.8 | 73 | Delta | 3 | 3 | 2 |
| AY.9 | 70 | Delta | 3 | 3 | 4 |
| BA.1.1.1 | 69 | Omicron | 4 | 2 | 5 |
| AY.36 | 68 | Delta | 3 | 3 | 2 |
| B.1.177.16 | 63 | Pre-VOC | 2 | 1 | 5 |
| B.1.201 | 63 | Pre-VOC | 1 | 2 | 4 |
| B | 61 | Pre-VOC | 1 | 2 | 5 |
| B.1.225 | 55 | Pre-VOC | 1 | 2 | 4 |
| BA.4 | 53 | Omicron | 5 | 2 | 2 |
| AY.46.5 | 52 | Delta | 3 | 3 | 4 |
| B.1.36.2 | 52 | Pre-VOC | 2 | 1 | 3 |
| AY.111 | 51 | Delta | 3 | 3 | 1 |
| AY.90 | 50 | Delta | 3 | 3 | 5 |
| AY.7 | 49 | Delta | 3 | 3 | 4 |
| B.1.258 | 49 | Pre-VOC | 2 | 3 | 5 |
| BQ.1.1 | 49 | Omicron | 7 | 3 | 5 |
| AY.122 | 47 | Delta | 3 | 3 | 1 |
| BQ.1 | 47 | Omicron | 7 | 3 | 5 |
| XL | 46 | Omicron | 5 | 2 | 5 |

| | | | | | |
|---|---|---|---|---|---|
| AY.9.2 | 43 | Delta | 3 | 3 | 5 |
| AD.2 | 42 | Pre-VOC | 2 | 3 | 1 |
| BE.1 | 42 | Omicron | 6 | 3 | 4 |
| BA.1.1.12 | 39 | Omicron | 4 | 2 | 5 |
| BA.2.3.5 | 37 | Omicron | 5 | 2 | 6 |
| XBB.1 | 37 | Omicron | 7 | 3 | 5 |
| AY.121 | 34 | Delta | 4 | 2 | 1 |
| AY.127 | 33 | Delta | 4 | 2 | 6 |
| BA.2.18 | 33 | Omicron | 5 | 2 | 2 |
| BQ.1.1.8 | 33 | Omicron | 8 | 2 | 5 |
| BA.2.12.1 | 31 | Omicron | 5 | 2 | 2 |
| BA.4.1 | 28 | Omicron | 6 | 3 | 2 |
| BE.1.1 | 28 | Omicron | 6 | 3 | 4 |
| AY.10 | 27 | Delta | 3 | 2 | 2 |
| BA.1.18 | 27 | Omicron | 4 | 2 | 5 |
| XE | 27 | Omicron | 5 | 2 | 5 |
| B.1.177.4 | 26 | Pre-VOC | 2 | 3 | 5 |
| BA.1.5 | 26 | Omicron | 4 | 2 | 5 |
| BF.28 | 26 | Omicron | 6 | 3 | 5 |
| BA.5.1.35 | 24 | Omicron | 6 | 2 | 3 |
| XBB.1.5.20 | 24 | Omicron | 8 | 2 | 5 |
| B.1.36.17 | 23 | Pre-VOC | 2 | 3 | 3 |
| AY.126 | 22 | Delta | 4 | 3 | 1 |
| B.1.177.10 | 22 | Pre-VOC | 2 | 3 | 5 |
| BA.2.23 | 22 | Omicron | 5 | 2 | 1 |
| BA.2.50 | 22 | Omicron | 5 | 2 | 2 |
| B.1.1.311 | 21 | Pre-VOC | 2 | 3 | 5 |
| B.1.1.37 | 21 | Pre-VOC | 2 | 3 | 5 |
| B.1.93 | 21 | Pre-VOC | 1 | 2 | 5 |
| BQ.1.1.62 | 21 | Omicron | 8 | 2 | 5 |
| XQ | 21 | Omicron | 5 | 2 | 5 |
| BA.1.10 | 20 | Omicron | 4 | 2 | 2 |
| BA.5.2.3 | 19 | Omicron | 6 | 3 | 4 |
| BA.5.2.6 | 19 | Omicron | 7 | 3 | 4 |
| BD.1 | 19 | Omicron | 4 | 2 | 3 |
| BF.11.2 | 19 | Omicron | 7 | 3 | 4 |
| BA.4.6 | 18 | Omicron | 6 | 3 | 3 |
| AY.4.11 | 17 | Delta | 4 | 2 | 6 |
| BQ.1.8 | 17 | Omicron | 7 | 3 | 5 |
| B.1.258.3 | 16 | Pre-VOC | 2 | 2 | 5 |
| BA.4.1.8 | 16 | Omicron | 6 | 3 | 3 |
| BA.5.1.30 | 16 | Omicron | 6 | 3 | 4 |
| BA.2.37 | 15 | Omicron | 5 | 2 | 2 |
| BA.5.1.33 | 15 | Omicron | 7 | 3 | 4 |
| B.1.160 | 14 | Pre-VOC | 2 | 3 | 2 |
| B.1.177.54 | 14 | Pre-VOC | 2 | 3 | 4 |
| B.1.177.60 | 14 | Pre-VOC | 2 | 1 | 2 |
| B.1.177.69 | 14 | Pre-VOC | 2 | 2 | 5 |
| BA.5.2.13 | 14 | Omicron | 7 | 3 | 3 |
| BA.1.7 | 13 | Omicron | 4 | 2 | 2 |
| BN.1.3 | 13 | Omicron | 8 | 2 | 4 |
| B.1.177.87 | 12 | Pre-VOC | 2 | 2 | 4 |
| B.1.36.9 | 12 | Pre-VOC | 2 | 3 | 4 |
| B.1.1.307 | 11 | Pre-VOC | 2 | 3 | 5 |
| BA.2.12 | 11 | Omicron | 5 | 2 | 2 |
| BA.2.8 | 11 | Omicron | 5 | 2 | 1 |
| BA.4.6.1 | 11 | Omicron | 7 | 3 | 3 |
| BA.5 | 11 | Omicron | 6 | 3 | 4 |
| BQ.1.1.1 | 11 | Omicron | 7 | 3 | 5 |
| B.1.1.3 | 10 | Pre-VOC | 1 | 2 | 5 |

| | | | | | |
|---|---|---|---|---|---|
| B.1.177.17 | 10 | Pre-VOC | 2 | 3 | 4 |
| B.1.221 | 10 | Pre-VOC | 2 | 1 | 5 |
| B.3 | 10 | Pre-VOC | 1 | 2 | 5 |
| B.40 | 10 | Pre-VOC | 1 | 2 | 5 |
| BA.5.1.10 | 10 | Omicron | 6 | 2 | 3 |
| BA.5.1.5 | 10 | Omicron | 6 | 3 | 3 |
| BA.5.2.26 | 10 | Omicron | 6 | 3 | 4 |
| BE.1.2 | 10 | Omicron | 7 | 3 | 3 |
| BF.4 | 10 | Omicron | 6 | 2 | 4 |
| BF.7 | 10 | Omicron | 7 | 3 | 4 |
| B.1.111 | 9 | Pre-VOC | 1 | 2 | 5 |
| BA.1.12 | 9 | Omicron | 4 | 2 | 5 |
| BA.5.1.21 | 9 | Omicron | 6 | 2 | 3 |
| BA.5.2.20 | 9 | Omicron | 6 | 3 | 3 |
| BA.5.2.7 | 9 | Omicron | 7 | 2 | 3 |
| BA.5.2.9 | 9 | Omicron | 6 | 3 | 4 |
| BA.5.3.3 | 9 | Omicron | 6 | 3 | 4 |
| BF.5 | 9 | Omicron | 6 | 2 | 5 |
| BF.6 | 9 | Omicron | 6 | 2 | 5 |
| BQ.1.1.4 | 9 | Omicron | 8 | 2 | 4 |
| AY.121.1 | 8 | Delta | 4 | 2 | 6 |
| AY.4.5 | 8 | Delta | 3 | 3 | 5 |
| AY.87 | 8 | Delta | 3 | 3 | 5 |
| B.1.351 | 8 | Beta | 2 | 1 | 4 |
| BA.5.1.22 | 8 | Omicron | 6 | 3 | 4 |
| BA.5.2.35 | 8 | Omicron | 7 | 3 | 4 |
| BQ.1.1.2 | 8 | Omicron | 7 | 3 | 5 |
| DF.1 | 8 | Omicron | 7 | 3 | 5 |
| AY.34 | 7 | Delta | 3 | 3 | 1 |
| AY.34.1 | 7 | Delta | 4 | 2 | 1 |
| B.1.177.56 | 7 | Pre-VOC | 2 | 3 | 5 |
| B.1.177.81 | 7 | Pre-VOC | 2 | 2 | 5 |
| BA.1.20 | 7 | Omicron | 4 | 2 | 5 |
| BA.2.16 | 7 | Omicron | 5 | 2 | 2 |
| BA.5.1.23 | 7 | Omicron | 6 | 3 | 4 |
| BA.5.8 | 7 | Omicron | 6 | 2 | 3 |
| BE.1.1.2 | 7 | Omicron | 6 | 2 | 4 |
| BF.1 | 7 | Omicron | 6 | 2 | 4 |
| BF.26 | 7 | Omicron | 7 | 3 | 5 |
| BF.7.5 | 7 | Omicron | 7 | 3 | 5 |
| BQ.1.1.22 | 7 | Omicron | 8 | 2 | 5 |
| AY.46.6 | 6 | Delta | 4 | 3 | 2 |
| AY.8 | 6 | Delta | 3 | 2 | 5 |
| B.1.1.351 | 6 | Pre-VOC | 2 | 2 | 5 |
| BA.1.1.18 | 6 | Omicron | 5 | 2 | 5 |
| BA.1.14 | 6 | Omicron | 4 | 2 | 5 |
| BA.5.9 | 6 | Omicron | 6 | 3 | 3 |
| BF.11.1 | 6 | Omicron | 6 | 3 | 4 |
| BN.1 | 6 | Omicron | 8 | 2 | 5 |
| BQ.1.11 | 6 | Omicron | 7 | 3 | 5 |
| C.35 | 6 | Pre-VOC | 2 | 3 | 5 |
| AY.103 | 5 | Delta | 3 | 3 | 1 |
| AY.42 | 5 | Delta | 3 | 3 | 5 |
| AY.98.1 | 5 | Delta | 3 | 3 | 5 |
| B.1.1.1 | 5 | Pre-VOC | 1 | 3 | 5 |
| B.1.1.10 | 5 | Pre-VOC | 1 | 2 | 4 |
| B.1.1.12 | 5 | Pre-VOC | 2 | 2 | 5 |
| B.1.1.170 | 5 | Pre-VOC | 2 | 2 | 5 |
| B.1.221.1 | 5 | Pre-VOC | 2 | 2 | 4 |
| B.1.391 | 5 | Pre-VOC | 1 | 2 | 3 |

| | | | | | |
|---|---|---|---|---|---|
| B.33 | 5 | Pre-VOC | 1 | 2 | 5 |
| BA.2.32 | 5 | Omicron | 5 | 2 | 2 |
| BA.2.36 | 5 | Omicron | 5 | 2 | 1 |
| BA.2.75.1 | 5 | Omicron | 6 | 2 | 2 |
| BA.4.4 | 5 | Omicron | 6 | 2 | 4 |
| BA.5.1.24 | 5 | Omicron | 6 | 2 | 3 |
| BA.5.2.21 | 5 | Omicron | 7 | 2 | 3 |
| BA.5.6 | 5 | Omicron | 6 | 3 | 3 |
| BF.14 | 5 | Omicron | 7 | 3 | 5 |
| BN.1.5 | 5 | Omicron | 7 | 3 | 5 |
| BQ.1.13 | 5 | Omicron | 7 | 2 | 5 |
| CR.2 | 5 | Omicron | 7 | 2 | 5 |
| AY.25.1 | 4 | Delta | 4 | 2 | 3 |
| B.1.177.18 | 4 | Pre-VOC | 2 | 2 | 5 |
| B.1.177.7 | 4 | Pre-VOC | 2 | 3 | 5 |
| B.23 | 4 | Pre-VOC | 1 | 2 | 5 |
| BA.1.1.4 | 4 | Omicron | 4 | 2 | 5 |
| BA.2.22 | 4 | Omicron | 5 | 2 | 6 |
| BA.2.23.1 | 4 | Omicron | 5 | 2 | 1 |
| BA.2.38 | 4 | Omicron | 6 | 2 | 1 |
| BA.2.41 | 4 | Omicron | 5 | 2 | 1 |
| BA.5.1.2 | 4 | Omicron | 6 | 2 | 4 |
| BA.5.1.25 | 4 | Omicron | 6 | 2 | 3 |
| BA.5.1.26 | 4 | Omicron | 7 | 2 | 3 |
| BA.5.1.3 | 4 | Omicron | 6 | 2 | 3 |
| BA.5.2.28 | 4 | Omicron | 7 | 2 | 4 |
| BF.11 | 4 | Omicron | 6 | 3 | 4 |
| BQ.1.1.6 | 4 | Omicron | 7 | 3 | 5 |
| BQ.1.2 | 4 | Omicron | 7 | 3 | 4 |
| CH.1.1 | 4 | Omicron | 8 | 2 | 5 |
| CK.2.1 | 4 | Omicron | 7 | 2 | 5 |
| CN.1 | 4 | Omicron | 7 | 3 | 5 |
| AY.118 | 3 | Delta | 4 | 2 | 3 |
| AY.20.1 | 3 | Delta | 4 | 2 | 6 |
| AY.4.10 | 3 | Delta | 4 | 2 | 1 |
| B.1.1.279 | 3 | Pre-VOC | 2 | 2 | 5 |
| B.1.258.4 | 3 | Pre-VOC | 2 | 2 | 4 |
| B.1.36.39 | 3 | Pre-VOC | 2 | 2 | 4 |
| BA.1.19 | 3 | Omicron | 4 | 2 | 5 |
| BA.2.10.1 | 3 | Omicron | 5 | 2 | 5 |
| BA.2.39 | 3 | Omicron | 5 | 2 | 2 |
| BA.2.56 | 3 | Omicron | 6 | 2 | 1 |
| BA.4.1.1 | 3 | Omicron | 6 | 2 | 3 |
| BA.5.2.2 | 3 | Omicron | 6 | 2 | 4 |
| BA.5.3.1 | 3 | Omicron | 6 | 2 | 4 |
| BF.10 | 3 | Omicron | 7 | 2 | 5 |
| BN.1.3.1 | 3 | Omicron | 8 | 2 | 4 |
| BQ.1.3.2 | 3 | Omicron | 8 | 2 | 5 |
| CK.2.1.1 | 3 | Omicron | 7 | 2 | 5 |
| XAZ | 3 | Omicron | 6 | 2 | 5 |
| AY.107 | 2 | Delta | 3 | 2 | 4 |
| AY.4.9 | 2 | Delta | 3 | 3 | 5 |
| AY.47 | 2 | Delta | 4 | 2 | 5 |
| AY.58 | 2 | Delta | 3 | 2 | 4 |
| B.1.1.41 | 2 | Pre-VOC | 1 | 2 | 5 |
| B.1.1.500 | 2 | Pre-VOC | 2 | 2 | 4 |
| B.1.1.529 | 2 | Omicron | 5 | 2 | 4 |
| B.1.177.19 | 2 | Pre-VOC | 2 | 3 | 5 |
| B.1.177.20 | 2 | Pre-VOC | 2 | 2 | 4 |
| B.1.177.58 | 2 | Pre-VOC | 2 | 2 | 5 |

| | | | | | |
|---|---|---|---|---|---|
| B.1.177.65 | 2 | Pre-VOC | 2 | 2 | 5 |
| B.1.2 | 2 | Pre-VOC | 2 | 2 | 4 |
| B.1.240 | 2 | Pre-VOC | 2 | 2 | 1 |
| B.1.250 | 2 | Pre-VOC | 1 | 2 | 3 |
| B.1.36.7 | 2 | Pre-VOC | 2 | 2 | 4 |
| B.1.367 | 2 | Pre-VOC | 2 | 2 | 4 |
| B.1.389 | 2 | Pre-VOC | 2 | 2 | 4 |
| B.1.408 | 2 | Pre-VOC | 2 | 2 | 4 |
| B.1.610 | 2 | Pre-VOC | 2 | 2 | 4 |
| B.29 | 2 | Pre-VOC | 1 | 2 | 5 |
| BA.1.1.10 | 2 | Omicron | 5 | 2 | 5 |
| BA.1.1.7 | 2 | Omicron | 4 | 2 | 2 |
| BA.1.21 | 2 | Omicron | 4 | 2 | 2 |
| BA.2.10.3 | 2 | Omicron | 5 | 2 | 2 |
| BA.2.2.1 | 2 | Omicron | 5 | 2 | 1 |
| BA.2.51 | 2 | Omicron | 5 | 2 | 1 |
| BA.2.6 | 2 | Omicron | 5 | 2 | 2 |
| BA.2.76 | 2 | Omicron | 6 | 2 | 6 |
| BA.2.9.1 | 2 | Omicron | 5 | 2 | 2 |
| BA.2.9.3 | 2 | Omicron | 6 | 2 | 1 |
| BA.4.1.10 | 2 | Omicron | 6 | 3 | 3 |
| BA.4.1.5 | 2 | Omicron | 6 | 2 | 3 |
| BA.4.7 | 2 | Omicron | 6 | 2 | 3 |
| BA.5.1.1 | 2 | Omicron | 6 | 2 | 4 |
| BA.5.2.33 | 2 | Omicron | 6 | 3 | 4 |
| BA.5.2.34 | 2 | Omicron | 8 | 2 | 4 |
| BA.5.3 | 2 | Omicron | 7 | 2 | 3 |
| BA.5.3.2 | 2 | Omicron | 6 | 2 | 4 |
| BA.5.5 | 2 | Omicron | 6 | 2 | 4 |
| BE.1.1.1 | 2 | Omicron | 6 | 2 | 4 |
| BE.1.3 | 2 | Omicron | 6 | 2 | 3 |
| BF.11.5 | 2 | Omicron | 7 | 2 | 4 |
| BF.36 | 2 | Omicron | 7 | 3 | 5 |
| BF.7.4 | 2 | Omicron | 7 | 2 | 4 |
| BF.7.4.1 | 2 | Omicron | 8 | 2 | 4 |
| BQ.1.1.19 | 2 | Omicron | 8 | 2 | 5 |
| BQ.1.1.24 | 2 | Omicron | 8 | 2 | 5 |
| BQ.1.1.5 | 2 | Omicron | 7 | 2 | 5 |
| BQ.1.1.7 | 2 | Omicron | 8 | 2 | 4 |
| BQ.1.14 | 2 | Omicron | 7 | 2 | 4 |
| BS.1 | 2 | Omicron | 7 | 2 | 5 |
| CH.1.1.2 | 2 | Omicron | 8 | 2 | 5 |
| CJ.1.2 | 2 | Omicron | 8 | 2 | 5 |
| EC.1.1 | 2 | Omicron | 8 | 2 | 5 |
| Q.1 | 2 | Alpha | 3 | 2 | 5 |
| XAY.1 | 2 | Omicron | 8 | 2 | 5 |
| XN | 2 | Omicron | 5 | 2 | 5 |
| AY.129 | 1 | Delta | 3 | 2 | 1 |
| AY.25 | 1 | Delta | 4 | 2 | 1 |
| AY.3 | 1 | Delta | 4 | 2 | 1 |
| AY.32 | 1 | Delta | 3 | 2 | 3 |
| AY.33 | 1 | Delta | 3 | 2 | 3 |
| AY.4.13 | 1 | Delta | 4 | 2 | 2 |
| AY.4.17 | 1 | Delta | 3 | 2 | 1 |
| AY.4.2.3 | 1 | Delta | 4 | 2 | 3 |
| AY.4.4 | 1 | Delta | 3 | 2 | 4 |
| AY.45 | 1 | Delta | 3 | 2 | 5 |
| AY.88 | 1 | Delta | 4 | 2 | 4 |
| AY.99 | 1 | Delta | 4 | 2 | 4 |
| B.1.1.147 | 1 | Pre-VOC | 1 | 2 | 5 |

| | | | | | |
|---|---|---|---|---|---|
| B.1.1.159 | 1 | Pre-VOC | 2 | 2 | 4 |
| B.1.1.164 | 1 | Pre-VOC | 1 | 2 | 5 |
| B.1.1.189 | 1 | Pre-VOC | 1 | 2 | 5 |
| B.1.1.196 | 1 | Pre-VOC | 2 | 2 | 5 |
| B.1.1.204 | 1 | Pre-VOC | 2 | 2 | 5 |
| B.1.1.218 | 1 | Pre-VOC | 2 | 2 | 5 |
| B.1.1.222 | 1 | Pre-VOC | 2 | 2 | 5 |
| B.1.1.240 | 1 | Pre-VOC | 2 | 2 | 5 |
| B.1.1.274 | 1 | Pre-VOC | 2 | 2 | 5 |
| B.1.1.294 | 1 | Pre-VOC | 2 | 2 | 4 |
| B.1.1.301 | 1 | Pre-VOC | 2 | 2 | 5 |
| B.1.1.341 | 1 | Pre-VOC | 2 | 2 | 5 |
| B.1.1.347 | 1 | Pre-VOC | 2 | 2 | 5 |
| B.1.1.371 | 1 | Pre-VOC | 2 | 2 | 5 |
| B.1.1.487 | 1 | Pre-VOC | 2 | 2 | 5 |
| B.1.1.89 | 1 | Pre-VOC | 1 | 2 | 5 |
| B.1.160.23 | 1 | Pre-VOC | 2 | 2 | 5 |
| B.1.160.7 | 1 | Pre-VOC | 2 | 2 | 5 |
| B.1.177.15 | 1 | Pre-VOC | 2 | 2 | 5 |
| B.1.177.2 | 1 | Pre-VOC | 2 | 2 | 5 |
| B.1.177.48 | 1 | Pre-VOC | 2 | 2 | 5 |
| B.1.177.5 | 1 | Pre-VOC | 2 | 2 | 4 |
| B.1.177.55 | 1 | Pre-VOC | 2 | 2 | 5 |
| B.1.177.6 | 1 | Pre-VOC | 2 | 2 | 4 |
| B.1.177.8 | 1 | Pre-VOC | 2 | 2 | 3 |
| B.1.218 | 1 | Pre-VOC | 2 | 2 | 4 |
| B.1.222 | 1 | Pre-VOC | 1 | 2 | 4 |
| B.1.235 | 1 | Pre-VOC | 2 | 2 | 4 |
| B.1.236 | 1 | Pre-VOC | 2 | 2 | 5 |
| B.1.243 | 1 | Pre-VOC | 2 | 2 | 4 |
| B.1.258.12 | 1 | Pre-VOC | 2 | 2 | 4 |
| B.1.258.17 | 1 | Pre-VOC | 3 | 2 | 4 |
| B.1.36 | 1 | Pre-VOC | 1 | 2 | 4 |
| B.1.416 | 1 | Pre-VOC | 2 | 2 | 4 |
| B.1.416.1 | 1 | Pre-VOC | 2 | 2 | 4 |
| B.1.465 | 1 | Pre-VOC | 1 | 2 | 4 |
| B.1.471 | 1 | Pre-VOC | 2 | 2 | 3 |
| B.1.523 | 1 | Pre-VOC | 2 | 2 | 4 |
| B.1.540 | 1 | Pre-VOC | 2 | 2 | 2 |
| B.1.76 | 1 | Pre-VOC | 1 | 2 | 5 |
| B.1.9.5 | 1 | Pre-VOC | 2 | 2 | 5 |
| B.27 | 1 | Pre-VOC | 1 | 2 | 5 |
| B.28 | 1 | Pre-VOC | 1 | 2 | 5 |
| B.3.1 | 1 | Pre-VOC | 1 | 2 | 5 |
| B.39 | 1 | Pre-VOC | 1 | 2 | 5 |
| B.52 | 1 | Pre-VOC | 1 | 2 | 5 |
| BA.1.1.11 | 1 | Omicron | 5 | 2 | 5 |
| BA.2.13 | 1 | Omicron | 6 | 2 | 2 |
| BA.2.14 | 1 | Omicron | 5 | 2 | 1 |
| BA.2.25 | 1 | Omicron | 5 | 2 | 6 |
| BA.2.3.2 | 1 | Omicron | 5 | 2 | 2 |
| BA.2.3.9 | 1 | Omicron | 5 | 2 | 1 |
| BA.2.40.1 | 1 | Omicron | 6 | 2 | 1 |
| BA.2.44 | 1 | Omicron | 5 | 2 | 2 |
| BA.2.45 | 1 | Omicron | 5 | 2 | 2 |
| BA.2.52 | 1 | Omicron | 5 | 2 | 1 |
| BA.2.75 | 1 | Omicron | 7 | 2 | 1 |
| BA.2.75.5 | 1 | Omicron | 8 | 2 | 1 |
| BA.4.6.5 | 1 | Omicron | 7 | 2 | 4 |
| BA.5.1.6 | 1 | Omicron | 7 | 2 | 3 |

| | | | | | |
|---|---|---|---|---|---|
| BA.5.1.7 | 1 | Omicron | 6 | 2 | 4 |
| BA.5.2.16 | 1 | Omicron | 6 | 2 | 4 |
| BA.5.2.56 | 1 | Omicron | 7 | 2 | 3 |
| BA.5.6.2 | 1 | Omicron | 7 | 2 | 3 |
| BE.1.4 | 1 | Omicron | 6 | 2 | 3 |
| BE.5 | 1 | Omicron | 6 | 2 | 3 |
| BF.21 | 1 | Omicron | 7 | 2 | 4 |
| BF.27 | 1 | Omicron | 6 | 2 | 5 |
| BF.39 | 1 | Omicron | 8 | 2 | 4 |
| BF.5.1 | 1 | Omicron | 8 | 2 | 5 |
| BF.5.5 | 1 | Omicron | 7 | 2 | 5 |
| BF.7.23 | 1 | Omicron | 7 | 2 | 5 |
| BF.7.24 | 1 | Omicron | 7 | 2 | 4 |
| BF.7.3 | 1 | Omicron | 7 | 2 | 5 |
| BF.7.6 | 1 | Omicron | 7 | 2 | 4 |
| BF.7.7 | 1 | Omicron | 7 | 2 | 4 |
| BF.8 | 1 | Omicron | 7 | 2 | 5 |
| BF.9 | 1 | Omicron | 7 | 2 | 5 |
| BL.2 | 1 | Omicron | 7 | 2 | 5 |
| BM.1.1.3 | 1 | Omicron | 8 | 2 | 4 |
| BN.1.2 | 1 | Omicron | 8 | 2 | 4 |
| BN.1.4 | 1 | Omicron | 8 | 2 | 5 |
| BN.1.7 | 1 | Omicron | 8 | 2 | 5 |
| BQ.1.1.18 | 1 | Omicron | 8 | 2 | 5 |
| BQ.1.10 | 1 | Omicron | 8 | 2 | 5 |
| BQ.1.23 | 1 | Omicron | 8 | 2 | 5 |
| BQ.1.4 | 1 | Omicron | 7 | 2 | 5 |
| BQ.1.5 | 1 | Omicron | 7 | 2 | 5 |
| BQ.1.8.2 | 1 | Omicron | 8 | 2 | 5 |
| BU.1 | 1 | Omicron | 8 | 2 | 5 |
| BY.1.2 | 1 | Omicron | 7 | 2 | 5 |
| C.3 | 1 | Pre-VOC | 1 | 2 | 5 |
| CH.1.1.11 | 1 | Omicron | 8 | 2 | 5 |
| CQ.1 | 1 | Omicron | 7 | 2 | 5 |
| CR.1 | 1 | Omicron | 8 | 2 | 5 |
| CS.1 | 1 | Omicron | 7 | 2 | 5 |
| DN.1 | 1 | Omicron | 8 | 2 | 5 |
| DT.1 | 1 | Omicron | 8 | 2 | 5 |
| DV.6 | 1 | Omicron | 8 | 2 | 5 |
| EC.1 | 1 | Omicron | 8 | 2 | 5 |
| EF.1 | 1 | Omicron | 8 | 2 | 5 |
| EF.1.1 | 1 | Omicron | 8 | 2 | 5 |
| EF.2 | 1 | Omicron | 8 | 2 | 5 |
| P.2 | 1 | Zeta | 2 | 2 | 5 |
| XBB.2 | 1 | Omicron | 8 | 2 | 5 |
| Y.1 | 1 | Pre-VOC | 2 | 2 | 5 |

*401 unique lobal lineages observed within Norfolk, UK March 2020 – December 2022, in descending order of appearance. Variant of concern (VOC) designation was identified through Scorpio, as part of Pangolin. Lineages which appeared in the early pandemic before VOC designation (naming conventions began in May 2021) are labelled 'Pre-VOC'. First lineage emergence temporal cluster refers to 1 of 8 identified time periods with significant numbers of emerging lineages. Longevity/outdegree cluster refers to the assigned correlation strength cluster in the regression analysis. Assigned walktrap community refers to the subgroup identification.*

**Supplementary Table 3. SARS-CoV-2 lineages present during the Pre-VOC wave**

| Lineage | Appearances (n) | Appearances (%) | VOC |
|---|---|---|---|
| B.1.177 | 508 | 19.34 | Pre-VOC |
| B.1.1.7 | 399 | 15.19 | Alpha |
| B.1.1 | 305 | 11.61 | Pre-VOC |
| B.1.1.198 | 267 | 10.16 | Pre-VOC |
| B.1.177.57 | 194 | 7.38 | Pre-VOC |
| B.1 | 152 | 5.79 | Pre-VOC |
| B.1.201 | 63 | 2.40 | Pre-VOC |
| B | 61 | 2.32 | Pre-VOC |
| B.1.177.16 | 58 | 2.21 | Pre-VOC |
| B.1.225 | 55 | 2.09 | Pre-VOC |
| B.1.258 | 49 | 1.87 | Pre-VOC |
| B.1.36.2 | 46 | 1.75 | Pre-VOC |
| AD.2 | 42 | 1.60 | Pre-VOC |
| B.1.177.4 | 26 | 0.99 | Pre-VOC |
| B.1.36.17 | 23 | 0.88 | Pre-VOC |
| B.1.177.10 | 22 | 0.84 | Pre-VOC |
| B.1.1.311 | 21 | 0.80 | Pre-VOC |
| B.1.1.37 | 21 | 0.80 | Pre-VOC |
| B.1.93 | 21 | 0.80 | Pre-VOC |
| B.1.258.3 | 16 | 0.61 | Pre-VOC |
| B.1.160 | 14 | 0.53 | Pre-VOC |
| B.1.177.54 | 14 | 0.53 | Pre-VOC |
| B.1.36.9 | 12 | 0.46 | Pre-VOC |
| B.1.1.307 | 11 | 0.42 | Pre-VOC |
| B.1.177.87 | 11 | 0.42 | Pre-VOC |
| B.1.1.3 | 10 | 0.38 | Pre-VOC |
| B.1.177.17 | 10 | 0.38 | Pre-VOC |
| B.3 | 10 | 0.38 | Pre-VOC |
| B.40 | 10 | 0.38 | Pre-VOC |
| B.1.111 | 9 | 0.34 | Pre-VOC |
| B.1.221 | 9 | 0.34 | Pre-VOC |
| B.1.177.56 | 7 | 0.27 | Pre-VOC |
| B.1.177.81 | 7 | 0.27 | Pre-VOC |
| B.1.1.351 | 6 | 0.23 | Pre-VOC |
| B.1.177.60 | 6 | 0.23 | Pre-VOC |
| C.35 | 6 | 0.23 | Pre-VOC |
| B.1.1.1 | 5 | 0.19 | Pre-VOC |
| B.1.1.10 | 5 | 0.19 | Pre-VOC |
| B.1.1.12 | 5 | 0.19 | Pre-VOC |
| B.1.221.1 | 5 | 0.19 | Pre-VOC |
| B.1.391 | 5 | 0.19 | Pre-VOC |
| B.33 | 5 | 0.19 | Pre-VOC |
| B.1.177.18 | 4 | 0.15 | Pre-VOC |

| | | | |
|---|---|---|---|
| B.23 | 4 | 0.15 | Pre-VOC |

*Global lineages appearing in Norfolk during the defined Pre-VOC wave dated from the week commencing 09/03/2020 to 21/12/2021. Lineages present in the focused dataset highlighted in grey. Number of appearances listed in descending order, with appearances (%) calculated against the total number of sequences observed during this time frame, 2,627.*

**Supplementary Table 4. SARS-CoV-2 lineages present during the Alpha wave**

| Lineages | Appearances (n) | Appearances (%) | VOC |
|---|---|---|---|
| B.1.1.7 | 2106 | 92.94 | Alpha |
| B.1.177 | 58 | 2.56 | Pre-VOC |
| B.1.177.69 | 12 | 0.53 | Pre-VOC |
| B.1.617.2 | 10 | 0.44 | Delta |
| AY.10 | 9 | 0.40 | Delta |
| AY.4 | 8 | 0.35 | Delta |
| B.1.177.60 | 8 | 0.35 | Pre-VOC |
| B.1.351 | 7 | 0.31 | Pre-VOC |
| B.1.36.2 | 6 | 0.26 | Pre-VOC |
| B.1.1.198 | 5 | 0.22 | Pre-VOC |
| B.1.177.16 | 5 | 0.22 | Pre-VOC |
| B.1.177.57 | 5 | 0.22 | Pre-VOC |
| AY.7 | 4 | 0.18 | Delta |
| B.1.1 | 3 | 0.13 | Pre-VOC |
| B.1 | 2 | 0.09 | Pre-VOC |
| B.1.1.170 | 2 | 0.09 | Pre-VOC |
| B.1.2 | 2 | 0.09 | Pre-VOC |
| B.1.36.7 | 2 | 0.09 | Pre-VOC |
| Q.1 | 2 | 0.09 | Alpha |
| AY.9 | 1 | 0.04 | Delta |
| B.1.1.159 | 1 | 0.04 | Pre-VOC |
| B.1.1.222 | 1 | 0.04 | Pre-VOC |
| B.1.177.7 | 1 | 0.04 | Pre-VOC |
| B.1.177.87 | 1 | 0.04 | Pre-VOC |
| B.1.221 | 1 | 0.04 | Pre-VOC |
| B.1.243 | 1 | 0.04 | Pre-VOC |
| B.1.258.17 | 1 | 0.04 | Pre-VOC |
| B.1.408 | 1 | 0.04 | Pre-VOC |
| BA.1.1.15 | 1 | 0.04 | Omicron |

*Global lineages appearing in Norfolk during the defined Alpha wave dated from the week commencing 28/12/2020 to 24/05/2021. Lineages present in the focused dataset highlighted in grey. Number of appearances listed in descending order, with appearances (%) calculated against the total number of sequences observed during this time frame, 2266.*

**Supplementary Table 5. SARS-CoV-2 lineages present during the Delta wave**

| Lineage | Appearances (n) | Appearances (%) | VOC |
|---|---|---|---|
| AY.4 | 6479 | 62.40 | Delta |
| AY.4.2 | 813 | 7.83 | Delta |
| AY.6 | 393 | 3.79 | Delta |
| AY.43 | 372 | 3.58 | Delta |
| AY.98 | 333 | 3.21 | Delta |
| AY.5 | 303 | 2.92 | Delta |
| AY.120 | 207 | 1.99 | Delta |
| AY.4.2.2 | 207 | 1.99 | Delta |
| BA.1 | 127 | 1.22 | Omicron |
| B.1.617.2 | 96 | 0.92 | Delta |
| AY.124 | 93 | 0.90 | Delta |
| BA.1.17.2 | 77 | 0.74 | Omicron |
| AY.4.2.1 | 76 | 0.73 | Delta |
| AY.4.8 | 72 | 0.69 | Delta |
| AY.9 | 69 | 0.66 | Delta |
| AY.36 | 67 | 0.65 | Delta |
| AY.46.5 | 52 | 0.50 | Delta |
| AY.111 | 50 | 0.48 | Delta |
| AY.90 | 49 | 0.47 | Delta |
| AY.7 | 45 | 0.43 | Delta |
| AY.122 | 44 | 0.42 | Delta |
| AY.9.2 | 43 | 0.41 | Delta |
| AY.121 | 31 | 0.30 | Delta |
| AY.127 | 31 | 0.30 | Delta |
| B.1.1.7 | 31 | 0.30 | Alpha |
| BA.1.1 | 23 | 0.22 | Omicron |
| BA.1.15.1 | 23 | 0.22 | Omicron |
| AY.126 | 21 | 0.20 | Delta |
| AY.10 | 18 | 0.17 | Delta |
| BA.1.15 | 17 | 0.16 | Omicron |
| AY.4.11 | 16 | 0.15 | Delta |
| AY.121.1 | 8 | 0.08 | Delta |
| AY.4.5 | 8 | 0.08 | Delta |
| AY.87 | 8 | 0.08 | Delta |
| AY.34 | 7 | 0.07 | Delta |
| AY.34.1 | 7 | 0.07 | Delta |
| AY.46.6 | 6 | 0.06 | Delta |
| AY.8 | 6 | 0.06 | Delta |
| AY.98.1 | 5 | 0.05 | Delta |
| BA.1.16 | 5 | 0.05 | Omicron |
| BA.1.17 | 5 | 0.05 | Omicron |
| AY.25.1 | 4 | 0.04 | Delta |
| AY.42 | 4 | 0.04 | Delta |

| | | | |
|---|---|---|---|
| AY.118 | 3 | 0.03 | Delta |
| AY.20.1 | 3 | 0.03 | Delta |
| AY.103 | 2 | 0.02 | Delta |
| AY.107 | 2 | 0.02 | Delta |
| AY.4.9 | 2 | 0.02 | Delta |
| AY.47 | 2 | 0.02 | Delta |
| AY.58 | 2 | 0.02 | Delta |
| BA.1.1.15 | 2 | 0.02 | Omicron |
| AY.129 | 1 | 0.01 | Delta |
| AY.25 | 1 | 0.01 | Delta |
| AY.3 | 1 | 0.01 | Delta |
| AY.32 | 1 | 0.01 | Delta |
| AY.33 | 1 | 0.01 | Delta |
| AY.4.10 | 1 | 0.01 | Delta |
| AY.4.17 | 1 | 0.01 | Delta |
| AY.4.2.3 | 1 | 0.01 | Delta |
| AY.4.4 | 1 | 0.01 | Delta |
| AY.45 | 1 | 0.01 | Delta |
| AY.88 | 1 | 0.01 | Delta |
| AY.99 | 1 | 0.01 | Delta |
| B.1.351 | 1 | 0.01 | Beta |
| BA.1.14 | 1 | 0.01 | Omicron |

*Global lineages appearing in Norfolk during the defined Delta wave dated from the week commencing 31/05/2021 to 13/12/2021. Lineages present in the focused dataset highlighted in grey. Number of appearances listed in descending order, with appearances (%) calculated against the total number of sequences observed during this time frame, 10,383.*

**Supplementary Table 6. SARS-CoV-2 lineages present during the Omicron wave**

| Lineage | Appearances (n) | Appearances (%) | VOC |
|---|---|---|---|
| BA.2 | 3823 | 27.06 | Omicron |
| BA.1.1 | 2563 | 18.14 | Omicron |
| BA.1 | 1657 | 11.73 | Omicron |
| BA.1.17.2 | 1003 | 7.10 | Omicron |
| BA.1.1.13 | 769 | 5.44 | Omicron |
| BA.2.9 | 318 | 2.25 | Omicron |
| BA.2.1 | 298 | 2.11 | Omicron |
| BA.1.15.1 | 271 | 1.92 | Omicron |
| BA.1.1.14 | 250 | 1.77 | Omicron |
| BA.5.1 | 193 | 1.37 | Omicron |
| BA.1.1.15 | 149 | 1.05 | Omicron |
| BA.2.3 | 141 | 1.00 | Omicron |
| BA.2.10 | 140 | 0.99 | Omicron |
| BA.5.2.1 | 138 | 0.98 | Omicron |
| BA.1.17 | 136 | 0.96 | Omicron |
| BA.5.2 | 132 | 0.93 | Omicron |
| BA.1.15 | 116 | 0.82 | Omicron |
| BA.2.5 | 116 | 0.82 | Omicron |
| BA.1.16 | 103 | 0.73 | Omicron |
| BA.1.1.1 | 69 | 0.49 | Omicron |
| AY.4 | 53 | 0.38 | Delta |
| BA.4 | 53 | 0.38 | Omicron |
| BQ.1.1 | 49 | 0.35 | Omicron |
| BQ.1 | 47 | 0.33 | Omicron |
| XL | 46 | 0.33 | Omicron |
| BE.1 | 42 | 0.30 | Omicron |
| BA.1.1.12 | 39 | 0.28 | Omicron |
| BA.2.3.5 | 37 | 0.26 | Omicron |
| XBB.1 | 37 | 0.26 | Omicron |
| AY.4.2 | 36 | 0.25 | Delta |
| BA.2.18 | 33 | 0.23 | Omicron |
| BQ.1.1.8 | 33 | 0.23 | Omicron |
| BA.2.12.1 | 31 | 0.22 | Omicron |
| BA.4.1 | 28 | 0.20 | Omicron |
| BE.1.1 | 28 | 0.20 | Omicron |
| BA.1.18 | 27 | 0.19 | Omicron |
| XE | 27 | 0.19 | Omicron |
| BA.1.5 | 26 | 0.18 | Omicron |
| BF.28 | 26 | 0.18 | Omicron |
| BA.5.1.35 | 24 | 0.17 | Omicron |
| XBB.1.5.20 | 24 | 0.17 | Omicron |
| BA.2.23 | 22 | 0.16 | Omicron |
| BA.2.50 | 22 | 0.16 | Omicron |
| BQ.1.1.62 | 21 | 0.15 | Omicron |

| | | | |
|---|---|---|---|
| XQ | 21 | 0.15 | Omicron |
| BA.1.10 | 20 | 0.14 | Omicron |
| BA.5.2.3 | 19 | 0.13 | Omicron |
| BA.5.2.6 | 19 | 0.13 | Omicron |
| BD.1 | 19 | 0.13 | Omicron |
| BF.11.2 | 19 | 0.13 | Omicron |
| BA.4.6 | 18 | 0.13 | Omicron |
| BQ.1.8 | 17 | 0.12 | Omicron |
| BA.4.1.8 | 16 | 0.11 | Omicron |
| BA.5.1.30 | 16 | 0.11 | Omicron |
| AY.4.2.2 | 15 | 0.11 | Delta |
| BA.2.37 | 15 | 0.11 | Omicron |
| BA.5.1.33 | 15 | 0.11 | Omicron |
| BA.5.2.13 | 14 | 0.10 | Omicron |
| BA.1.7 | 13 | 0.09 | Omicron |
| BN.1.3 | 13 | 0.09 | Omicron |
| AY.43 | 11 | 0.08 | Delta |
| BA.2.12 | 11 | 0.08 | Omicron |
| BA.2.8 | 11 | 0.08 | Omicron |
| BA.4.6.1 | 11 | 0.08 | Omicron |
| BA.5 | 11 | 0.08 | Omicron |
| BQ.1.1.1 | 11 | 0.08 | Omicron |
| BA.5.1.10 | 10 | 0.07 | Omicron |
| BA.5.1.5 | 10 | 0.07 | Omicron |
| BA.5.2.26 | 10 | 0.07 | Omicron |
| BE.1.2 | 10 | 0.07 | Omicron |
| BF.4 | 10 | 0.07 | Omicron |
| BF.7 | 10 | 0.07 | Omicron |
| BA.1.12 | 9 | 0.06 | Omicron |
| BA.5.1.21 | 9 | 0.06 | Omicron |
| BA.5.2.20 | 9 | 0.06 | Omicron |
| BA.5.2.7 | 9 | 0.06 | Omicron |
| BA.5.2.9 | 9 | 0.06 | Omicron |
| BA.5.3.3 | 9 | 0.06 | Omicron |
| BF.5 | 9 | 0.06 | Omicron |
| BF.6 | 9 | 0.06 | Omicron |
| BQ.1.1.4 | 9 | 0.06 | Omicron |
| AY.98 | 8 | 0.06 | Delta |
| BA.5.1.22 | 8 | 0.06 | Omicron |
| BA.5.2.35 | 8 | 0.06 | Omicron |
| BQ.1.1.2 | 8 | 0.06 | Omicron |
| DF.1 | 8 | 0.06 | Omicron |
| BA.1.20 | 7 | 0.05 | Omicron |
| BA.2.16 | 7 | 0.05 | Omicron |
| BA.5.1.23 | 7 | 0.05 | Omicron |
| BA.5.8 | 7 | 0.05 | Omicron |

| | | | |
|---|---|---|---|
| BE.1.1.2 | 7 | 0.05 | Omicron |
| BF.1 | 7 | 0.05 | Omicron |
| BF.26 | 7 | 0.05 | Omicron |
| BF.7.5 | 7 | 0.05 | Omicron |
| BQ.1.1.22 | 7 | 0.05 | Omicron |
| AY.120 | 6 | 0.04 | Delta |
| BA.1.1.18 | 6 | 0.04 | Omicron |
| BA.5.9 | 6 | 0.04 | Omicron |
| BF.11.1 | 6 | 0.04 | Omicron |
| BN.1 | 6 | 0.04 | Omicron |
| BQ.1.11 | 6 | 0.04 | Omicron |
| BA.1.14 | 5 | 0.04 | Omicron |
| BA.2.32 | 5 | 0.04 | Omicron |
| BA.2.36 | 5 | 0.04 | Omicron |
| BA.2.75.1 | 5 | 0.04 | Omicron |
| BA.4.4 | 5 | 0.04 | Omicron |
| BA.5.1.24 | 5 | 0.04 | Omicron |
| BA.5.2.21 | 5 | 0.04 | Omicron |
| BA.5.6 | 5 | 0.04 | Omicron |
| BF.14 | 5 | 0.04 | Omicron |
| BN.1.5 | 5 | 0.04 | Omicron |
| BQ.1.13 | 5 | 0.04 | Omicron |
| CR.2 | 5 | 0.04 | Omicron |
| BA.1.1.4 | 4 | 0.03 | Omicron |
| BA.2.22 | 4 | 0.03 | Omicron |
| BA.2.23.1 | 4 | 0.03 | Omicron |
| BA.2.38 | 4 | 0.03 | Omicron |
| BA.2.41 | 4 | 0.03 | Omicron |
| BA.5.1.2 | 4 | 0.03 | Omicron |
| BA.5.1.25 | 4 | 0.03 | Omicron |
| BA.5.1.26 | 4 | 0.03 | Omicron |
| BA.5.1.3 | 4 | 0.03 | Omicron |
| BA.5.2.28 | 4 | 0.03 | Omicron |
| BF.11 | 4 | 0.03 | Omicron |
| BQ.1.1.6 | 4 | 0.03 | Omicron |
| BQ.1.2 | 4 | 0.03 | Omicron |
| CH.1.1 | 4 | 0.03 | Omicron |
| CK.2.1 | 4 | 0.03 | Omicron |
| CN.1 | 4 | 0.03 | Omicron |
| AY.103 | 3 | 0.02 | Delta |
| AY.121 | 3 | 0.02 | Delta |
| AY.122 | 3 | 0.02 | Delta |
| AY.4.2.1 | 3 | 0.02 | Delta |
| B.1.617.2 | 3 | 0.02 | Delta |
| BA.1.19 | 3 | 0.02 | Omicron |
| BA.2.10.1 | 3 | 0.02 | Omicron |

| | | | |
|---|---|---|---|
| BA.2.39 | 3 | 0.02 | Omicron |
| BA.2.56 | 3 | 0.02 | Omicron |
| BA.4.1.1 | 3 | 0.02 | Omicron |
| BA.5.2.2 | 3 | 0.02 | Omicron |
| BA.5.3.1 | 3 | 0.02 | Omicron |
| BF.10 | 3 | 0.02 | Omicron |
| BN.1.3.1 | 3 | 0.02 | Omicron |
| BQ.1.3.2 | 3 | 0.02 | Omicron |
| CK.2.1.1 | 3 | 0.02 | Omicron |
| XAZ | 3 | 0.02 | Omicron |
| AY.127 | 2 | 0.01 | Delta |
| AY.4.10 | 2 | 0.01 | Delta |
| B.1.1.529 | 2 | 0.01 | Omicron |
| BA.1.1.10 | 2 | 0.01 | Omicron |
| BA.1.1.7 | 2 | 0.01 | Omicron |
| BA.1.21 | 2 | 0.01 | Omicron |
| BA.2.10.3 | 2 | 0.01 | Omicron |
| BA.2.2.1 | 2 | 0.01 | Omicron |
| BA.2.51 | 2 | 0.01 | Omicron |
| BA.2.6 | 2 | 0.01 | Omicron |
| BA.2.76 | 2 | 0.01 | Omicron |
| BA.2.9.1 | 2 | 0.01 | Omicron |
| BA.2.9.3 | 2 | 0.01 | Omicron |
| BA.4.1.10 | 2 | 0.01 | Omicron |
| BA.4.1.5 | 2 | 0.01 | Omicron |
| BA.4.7 | 2 | 0.01 | Omicron |
| BA.5.1.1 | 2 | 0.01 | Omicron |
| BA.5.2.33 | 2 | 0.01 | Omicron |
| BA.5.2.34 | 2 | 0.01 | Omicron |
| BA.5.3 | 2 | 0.01 | Omicron |
| BA.5.3.2 | 2 | 0.01 | Omicron |
| BA.5.5 | 2 | 0.01 | Omicron |
| BE.1.1.1 | 2 | 0.01 | Omicron |
| BE.1.3 | 2 | 0.01 | Omicron |
| BF.11.5 | 2 | 0.01 | Omicron |
| BF.36 | 2 | 0.01 | Omicron |
| BF.7.4 | 2 | 0.01 | Omicron |
| BF.7.4.1 | 2 | 0.01 | Omicron |
| BQ.1.1.19 | 2 | 0.01 | Omicron |
| BQ.1.1.24 | 2 | 0.01 | Omicron |
| BQ.1.1.5 | 2 | 0.01 | Omicron |
| BQ.1.1.7 | 2 | 0.01 | Omicron |
| BQ.1.14 | 2 | 0.01 | Omicron |
| BS.1 | 2 | 0.01 | Omicron |
| CH.1.1.2 | 2 | 0.01 | Omicron |
| CJ.1.2 | 2 | 0.01 | Omicron |

| | | | |
|---|---|---|---|
| EC.1.1 | 2 | 0.01 | Omicron |
| XAY.1 | 2 | 0.01 | Omicron |
| XN | 2 | 0.01 | Omicron |
| AY.111 | 1 | 0.01 | Delta |
| AY.126 | 1 | 0.01 | Delta |
| AY.36 | 1 | 0.01 | Delta |
| AY.4.11 | 1 | 0.01 | Delta |
| AY.4.13 | 1 | 0.01 | Delta |
| AY.4.8 | 1 | 0.01 | Delta |
| AY.42 | 1 | 0.01 | Delta |
| AY.5 | 1 | 0.01 | Delta |
| AY.90 | 1 | 0.01 | Delta |
| BA.1.1.11 | 1 | 0.01 | Omicron |
| BA.2.13 | 1 | 0.01 | Omicron |
| BA.2.14 | 1 | 0.01 | Omicron |
| BA.2.25 | 1 | 0.01 | Omicron |
| BA.2.3.2 | 1 | 0.01 | Omicron |
| BA.2.3.9 | 1 | 0.01 | Omicron |
| BA.2.40.1 | 1 | 0.01 | Omicron |
| BA.2.44 | 1 | 0.01 | Omicron |
| BA.2.45 | 1 | 0.01 | Omicron |
| BA.2.52 | 1 | 0.01 | Omicron |
| BA.2.75 | 1 | 0.01 | Omicron |
| BA.2.75.5 | 1 | 0.01 | Omicron |
| BA.4.6.5 | 1 | 0.01 | Omicron |
| BA.5.1.6 | 1 | 0.01 | Omicron |
| BA.5.1.7 | 1 | 0.01 | Omicron |
| BA.5.2.16 | 1 | 0.01 | Omicron |
| BA.5.2.56 | 1 | 0.01 | Omicron |
| BA.5.6.2 | 1 | 0.01 | Omicron |
| BE.1.4 | 1 | 0.01 | Omicron |
| BE.5 | 1 | 0.01 | Omicron |
| BF.21 | 1 | 0.01 | Omicron |
| BF.27 | 1 | 0.01 | Omicron |
| BF.39 | 1 | 0.01 | Omicron |
| BF.5.1 | 1 | 0.01 | Omicron |
| BF.5.5 | 1 | 0.01 | Omicron |
| BF.7.23 | 1 | 0.01 | Omicron |
| BF.7.24 | 1 | 0.01 | Omicron |
| BF.7.3 | 1 | 0.01 | Omicron |
| BF.7.6 | 1 | 0.01 | Omicron |
| BF.7.7 | 1 | 0.01 | Omicron |
| BF.8 | 1 | 0.01 | Omicron |
| BF.9 | 1 | 0.01 | Omicron |
| BL.2 | 1 | 0.01 | Omicron |
| BM.1.1.3 | 1 | 0.01 | Omicron |

| | | | |
|---|---|---|---|
| BN.1.2 | 1 | 0.01 | Omicron |
| BN.1.4 | 1 | 0.01 | Omicron |
| BN.1.7 | 1 | 0.01 | Omicron |
| BQ.1.1.18 | 1 | 0.01 | Omicron |
| BQ.1.10 | 1 | 0.01 | Omicron |
| BQ.1.23 | 1 | 0.01 | Omicron |
| BQ.1.4 | 1 | 0.01 | Omicron |
| BQ.1.5 | 1 | 0.01 | Omicron |
| BQ.1.8.2 | 1 | 0.01 | Omicron |
| BU.1 | 1 | 0.01 | Omicron |
| BY.1.2 | 1 | 0.01 | Omicron |
| CH.1.1.11 | 1 | 0.01 | Omicron |
| CQ.1 | 1 | 0.01 | Omicron |
| CR.1 | 1 | 0.01 | Omicron |
| CS.1 | 1 | 0.01 | Omicron |
| DN.1 | 1 | 0.01 | Omicron |
| DT.1 | 1 | 0.01 | Omicron |
| DV.6 | 1 | 0.01 | Omicron |
| EC.1 | 1 | 0.01 | Omicron |
| EF.1 | 1 | 0.01 | Omicron |
| EF.1.1 | 1 | 0.01 | Omicron |
| EF.2 | 1 | 0.01 | Omicron |
| XBB.2 | 1 | 0.01 | Omicron |

*Global lineages appearing in Norfolk during the defined Omicron wave dated from the week commencing 20/12/2021 to 26/12/2022. Lineages present in the focused dataset highlighted in grey. Number of appearances listed in descending order, with appearances (%) calculated against the total number of sequences observed during this time frame, 14,130.*

**Supplementary Table 7. Top appearing lineages by study counties**

| | Norfolk | | Suffolk | | Hertfordshire | |
|---|---|---|---|---|---|---|
| | Lineage | % Appearance | Lineage | % Appearance | Lineage | % Appearance |
| **Pre-VOC** | B.1.177 | 19.3 | B.1.177 | 23.8 | B.1.1 | 22.4 |
| | B.1.1.7 | 15.2 | B.1.1 | 13.1 | B.1.177 | 12.3 |
| | B.1.1 | 11.6 | B.1 | 8.8 | B.1 | 7.4 |
| **Alpha** | B.1.1.7 | 92.9 | B.1.1.7 | 88.9 | B.1.1.7 | 87.7 |
| | B.1.177 | 2.6 | B.1.177 | 6.3 | B.1.177 | 2.9 |
| | B.1.177.69 | 0.5 | AY.4 | 0.9 | B.1.617.2 | 1.2 |
| **Delta** | AY.4 | 62.4 | AY.4 | 59.6 | AY.4 | 59.2 |
| | AY.4.2 | 7.8 | AY.98 | 4.9 | AY.4.2 | 8.0 |
| | AY.6 | 3.8 | AY.4.2 | 4.8 | AY.43 | 4.5 |
| **Omicron** | BA.2 | 30.5 | BA.2 | 31.7 | BA.2 | 27.6 |
| | BA.1.1 | 18.1 | BA.1.1 | 16.7 | BA.1.1 | 14.8 |
| | BA.1 | 11.7 | BA.1 | 13.7 | BA.1 | 14.8 |

*Most frequently appearing SARS-CoV-2 lineages in each dated wave in Norfolk and comparison counties Suffolk and Hertfordshire. % Appearance refers to the overall number of sequences present of the respective lineage within the wave.*
