## Supplementary material for "Genomic Epidemiology of SARS-CoV-2 in Norfolk, UK, March 2020 – December 2022": COG-UK Consortium Author List

**Funding acquisition, Leadership and supervision, Metadata curation, Project administration, Samples and logistics, Sequencing and analysis, Software and analysis tools, and Visualisation:**
Dr Samuel C Robson PhD [13, 84]

**Funding acquisition, Leadership and supervision, Metadata curation, Project administration, Samples and logistics, Sequencing and analysis, and Software and analysis tools:**
Dr Thomas R Connor PhD [11, 74] and Prof Nicholas J Loman PhD [43]

**Leadership and supervision, Metadata curation, Project administration, Samples and logistics, Sequencing and analysis, Software and analysis tools, and Visualisation:**
Dr Tanya Golubchik PhD [5]

**Funding acquisition, Leadership and supervision, Metadata curation, Samples and logistics, Sequencing and analysis, and Visualisation:**
Dr Rocio T Martinez Nunez PhD [46]

**Funding acquisition, Leadership and supervision, Project administration, Samples and logistics, Sequencing and analysis, and Software and analysis tools:**
Dr David Bonsall PhD [5]

**Funding acquisition, Leadership and supervision, Project administration, Sequencing and analysis, Software and analysis tools, and Visualisation:**
Prof Andrew Rambaut DPhil [104]

**Funding acquisition, Metadata curation, Project administration, Samples and logistics, Sequencing and analysis, and Software and analysis tools:**
Dr Luke B Snell MSc, MBBS [12]

**Leadership and supervision, Metadata curation, Project administration, Samples and logistics, Software and analysis tools, and Visualisation:**
Rich Livett MSc [116]

**Funding acquisition, Leadership and supervision, Metadata curation, Project administration, and Samples and logistics:**
Dr Catherine Ludden PhD [20, 70]

**Funding acquisition, Leadership and supervision, Metadata curation, Samples and logistics, and Sequencing and analysis:**
Dr Sally Corden PhD [74] and Dr Eleni Nastouli FRCPath [96, 95, 30]

**Funding acquisition, Leadership and supervision, Metadata curation, Sequencing and analysis, and Software and analysis tools:**
Dr Gaia Nebbia PhD, FRCPath [12]

**Funding acquisition, Leadership and supervision, Project administration, Samples and logistics, and Sequencing and analysis:**
Ian Johnston BSc [116]

**Leadership and supervision, Metadata curation, Project administration, Samples and logistics, and Sequencing and analysis:**
Prof Katrina Lythgoe PhD [5], Dr M. Estee Torok FRCP [19, 20] and Prof Ian G Goodfellow PhD [24]

**Leadership and supervision, Metadata curation, Project administration, Samples and logistics, and Visualisation:**
Dr Jacqui A Prieto PhD [97, 82] and Dr Kordo Saeed MD, FRCPath [97, 83]

**Leadership and supervision, Metadata curation, Project administration, Sequencing and analysis, and Software and analysis tools:**
Dr David K Jackson PhD [116]

**Leadership and supervision, Metadata curation, Samples and logistics, Sequencing and analysis, and Visualisation:**
Dr Catherine Houlihan PhD [96, 94]

**Leadership and supervision, Metadata curation, Sequencing and analysis, Software and analysis tools, and Visualisation:**
Dr Dan Frampton PhD [94, 95]

**Metadata curation, Project administration, Samples and logistics, Sequencing and analysis, and Software and analysis tools:**
Dr William L Hamilton PhD [19] and Dr Adam A Witney PhD [41]

**Funding acquisition, Samples and logistics, Sequencing and analysis, and Visualisation:**
Dr Giselda Bucca PhD [101]

**Funding acquisition, Leadership and supervision, Metadata curation, and Project administration:**
Dr Cassie F Pope PhD[40, 41]

**Funding acquisition, Leadership and supervision, Metadata curation, and Samples and logistics:**
Dr Catherine Moore PhD [74]

**Funding acquisition, Leadership and supervision, Metadata curation, and Sequencing and analysis:**
Prof Emma C Thomson PhD, FRCP [53]

**Funding acquisition, Leadership and supervision, Project administration, and Samples and logistics:**
Dr Teresa Cutino-Moguel PhD [2], Dr Ewan M Harrison PhD [116, 102]

**Funding acquisition, Leadership and supervision, Sequencing and analysis, and Visualisation:**
Prof Colin P Smith PhD [101]

**Leadership and supervision, Metadata curation, Project administration, and Sequencing and analysis:**
Fiona Rogan BSc [77]

**Leadership and supervision, Metadata curation, Project administration, and Samples and logistics:**
Shaun M Beckwith MSc [6], Abigail Murray Degree [6], Dawn Singleton HNC [6], Dr Kirstine Eastick PhD, FRCPath [37], Dr Liz A Sheridan PhD [98], Paul Randell MSc, PgD [99], Dr Leigh M Jackson PhD [105], Dr Cristina V Ariani PhD [116] and Dr Sónia Gonçalves PhD [116]

**Leadership and supervision, Metadata curation, Samples and logistics, and Sequencing and analysis:**
Dr Derek J Fairley PhD [3, 77], Prof Matthew W Loose PhD [18] and Joanne Watkins MSc [74]

**Leadership and supervision, Metadata curation, Samples and logistics, and Visualisation:**
Dr Samuel Moses MD [25, 106]

**Leadership and supervision, Metadata curation, Sequencing and analysis, and Software and analysis tools:**
Dr Sam Nicholls PhD [43], Dr Matthew Bull PhD [74] and Dr Roberto Amato PhD [116]

**Leadership and supervision, Project administration, Samples and logistics, and Sequencing and analysis:**
Prof Darren L Smith PhD [36, 65, 66]

**Leadership and supervision, Sequencing and analysis, Software and analysis tools, and Visualisation:**
Prof David M Aanensen PhD [14, 116] and Dr Jeffrey C Barrett PhD [116]

**Metadata curation, Project administration, Samples and logistics, and Sequencing and analysis:**
Dr Beatrix Kele PhD [2], Dr Dinesh Aggarwal MRCP[20, 116, 70], Dr James G Shepherd MBCHB, MRCP [53], Dr Martin D Curran PhD [71] and Dr Surendra Parmar PhD [71]

**Metadata curation, Project administration, Sequencing and analysis, and Software and analysis tools:**
Dr Matthew D Parker PhD [109]

**Metadata curation, Samples and logistics, Sequencing and analysis, and Software and analysis tools:**
Dr Catryn Williams PhD [74]

**Metadata curation, Samples and logistics, Sequencing and analysis, and Visualisation:**
Dr Sharon Glaysher PhD [68]

**Metadata curation, Sequencing and analysis, Software and analysis tools, and Visualisation:**
Dr Anthony P Underwood PhD [14, 116], Dr Matthew Bashton PhD [36, 65], Dr Nicole Pacchiarini PhD [74], Dr Katie F Loveson PhD [84] and Matthew Byott MSc [95, 96]

**Project administration, Sequencing and analysis, Software and analysis tools, and Visualisation:**
Dr Alessandro M Carabelli PhD [20]

**Funding acquisition, Leadership and supervision, and Metadata curation:**
Dr Kate E Templeton PhD [56, 104]

**Funding acquisition, Leadership and supervision, and Project administration:**
Prof Sharon J Peacock PhD [20, 70], Dr Thushan I de Silva PhD [109], Dr Dennis Wang PhD [109], Dr Cordelia F Langford PhD [116] and John Sillitoe BEng [116]

**Funding acquisition, Leadership and supervision, and Samples and logistics:**
Prof Rory N Gunson PhD, FRCPath [55]

**Funding acquisition, Leadership and supervision, and Sequencing and analysis:**
Dr Simon Cottrell PhD [74], Dr Justin O'Grady PhD [75, 103] and Prof Dominic Kwiatkowski PhD [116, 108]

**Leadership and supervision, Metadata curation, and Project administration:**
Dr Patrick J Lillie PhD, FRCP [37]

**Leadership and supervision, Metadata curation, and Samples and logistics:**
Dr Nicholas Cortes MBCHB [33], Dr Nathan Moore MBCHB [33], Dr Claire Thomas DPhil [33], Phillipa J Burns MSc, DipRCPath [37], Dr Tabitha W Mahungu FRCPath [80] and Steven Liggett BSc [86]

**Leadership and supervision, Metadata curation, and Sequencing and analysis:**
Angela H Beckett MSc [13, 81] and Prof Matthew TG Holden PhD [73]

**Leadership and supervision, Project administration, and Samples and logistics:**
Dr Lisa J Levett PhD [34], Dr Husam Osman PhD [70, 35] and Dr Mohammed O Hassan-Ibrahim PhD, FRCPath [99]

**Leadership and supervision, Project administration, and Sequencing and analysis:**
Dr David A Simpson PhD [77]

**Leadership and supervision, Samples and logistics, and Sequencing and analysis:**
Dr Meera Chand PhD [72], Prof Ravi K Gupta PhD [102], Prof Alistair C Darby PhD [107] and Prof Steve Paterson PhD [107]

**Leadership and supervision, Sequencing and analysis, and Software and analysis tools:**
Prof Oliver G Pybus DPhil [23], Dr Erik M Volz PhD [39], Prof Daniela de Angelis PhD [52], Prof David L Robertson PhD [53], Dr Andrew J Page PhD [75] and Dr Inigo Martincorena PhD [116]

**Leadership and supervision, Sequencing and analysis, and Visualisation:**
Dr Louise Aigrain PhD [116] and Dr Andrew R Bassett PhD [116]

**Metadata curation, Project administration, and Samples and logistics:**
Dr Nick Wong DPhil, MRCP, FRCPath [50], Dr Yusri Taha MD, PhD [89], Michelle J Erkiert BA [99] and Dr Michael H Spencer Chapman MBBS [116, 102]

**Metadata curation, Project administration, and Sequencing and analysis:**
Dr Rebecca Dewar PhD [56] and Martin P McHugh MSc [56, 111]

**Metadata curation, Project administration, and Software and analysis tools:**
Siddharth Mookerjee MPH [38, 57]

**Metadata curation, Project administration, and Visualisation:**
Stephen Aplin [97], Matthew Harvey [97], Thea Sass [97], Dr Helen Umpleby FRCP [97] and Helen Wheeler [97]

**Metadata curation, Samples and logistics, and Sequencing and analysis:**
Dr James P McKenna PhD [3], Dr Ben Warne MRCP [9], Joshua F Taylor MSc [22], Yasmin Chaudhry BSc [24], Rhys Izuagbe [24], Dr Aminu S Jahun PhD [24], Dr Gregory R Young PhD [36, 65], Dr Claire McMurray PhD [43], Dr Clare M McCann PhD [65, 66], Dr Andrew Nelson PhD [65, 66] and Scott Elliott [68]

**Metadata curation, Samples and logistics, and Visualisation:**
Hannah Lowe MSc [25]

**Metadata curation, Sequencing and analysis, and Software and analysis tools:**
Dr Anna Price PhD [11], Matthew R Crown BSc [65], Dr Sara Rey PhD [74], Dr Sunando Roy PhD [96] and Dr Ben Temperton PhD [105]

**Metadata curation, Sequencing and analysis, and Visualisation:**
Dr Sharif Shaaban PhD [73] and Dr Andrew R Hesketh PhD [101]

**Project administration, Samples and logistics, and Sequencing and analysis:**
Dr Kenneth G Laing PhD[41], Dr Irene M Monahan PhD [41] and Dr Judith Heaney PhD [95, 96, 34]

**Project administration, Samples and logistics, and Visualisation:**
Dr Emanuela Pelosi FRCPath [97], Siona Silviera MSc [97] and Dr Eleri Wilson-Davies MD,

FRCPath [97]

**Samples and logistics, Software and analysis tools, and Visualisation:**
Dr Helen Fryer PhD [5]

**Sequencing and analysis, Software and analysis tools, and Visualization:**
Dr Helen Adams PhD [4], Dr Louis du Plessis PhD [23], Dr Rob Johnson PhD [39], Dr William T Harvey PhD [53, 42], Dr Joseph Hughes PhD [53], Dr Richard J Orton PhD [53], Dr Lewis G Spurgin PhD [59], Dr Yann Bourgeois PhD [81], Dr Chris Ruis PhD [102], Áine O'Toole MSc [104], Marina Gourtovaia MSc [116] and Dr Theo Sanderson PhD [116]

**Funding acquisition, and Leadership and supervision:**
Dr Christophe Fraser PhD [5], Dr Jonathan Edgeworth PhD, FRCPath [12], Prof Judith Breuer MD [96, 29], Dr Stephen L Michell PhD [105] and Prof John A Todd PhD [115]

**Funding acquisition, and Project administration:**
Michaela John BSc [10] and Dr David Buck PhD [115]

**Leadership and supervision, and Metadata curation:**
Dr Kavitha Gajee MBBS, FRCPath [37] and Dr Gemma L Kay PhD [75]

**Leadership and supervision, and Project administration:**
David Heyburn [74]

**Leadership and supervision, and Samples and logistics:**
Dr Themoula Charalampous PhD [12, 46], Adela Alcolea-Medina [32, 112], Katie Kitchman BSc [37], Prof Alan McNally PhD [43, 93], David T Pritchard MSc, CSci [50], Dr Samir Dervisevic FRCPath [58], Dr Peter Muir PhD [70], Dr Esther Robinson PhD [70, 35], Dr Barry B Vipond PhD [70], Newara A Ramadan MSc, CSci, FIBMS [78], Dr Christopher Jeanes MBBS [90], Danni Weldon BSc [116], Jana Catalan MSc [118] and Neil Jones MSc [118]

**Leadership and supervision, and Sequencing and analysis:**
Dr Ana da Silva Filipe PhD [53], Dr Chris Williams MBBS [74], Marc Fuchs BSc [77], Dr Julia Miskelly PhD [77], Dr Aaron R Jeffries PhD [105], Karen Oliver BSc [116] and Dr Naomi R Park PhD [116]

**Metadata curation, and Samples and logistics:**
Amy Ash BSc [1], Cherian Koshy MSc, CSci, FIBMS [1], Magdalena Barrow [7], Dr Sarah L Buchan PhD [7], Dr Anna Mantzouratou PhD [7], Dr Gemma Clark PhD [15], Dr Christopher W Holmes PhD [16], Sharon Campbell MSc [17], Thomas Davis MSc [21], Ngee Keong Tan MSc [22], Dr Julianne R Brown PhD [29], Dr Kathryn A Harris PhD [29, 2], Stephen P Kidd MSc [33], Dr Paul R Grant PhD [34], Dr Li Xu-McCrae PhD [35], Dr Alison Cox PhD [38, 63], Pinglawathee Madona [38, 63], Dr Marcus Pond PhD [38, 63], Dr Paul A Randell MBBCh [38, 63], Karen T Withell FIBMS [48], Cheryl Williams MSc [51], Dr Clive Graham MD [60], Rebecca Denton-Smith BSc [62], Emma Swindells BSc [62], Robyn Turnbull BSc [62], Dr Tim J Sloan PhD [67], Dr Andrew Bosworth PhD [70, 35], Stephanie Hutchings [70], Hannah M Pymont MSc [70], Dr Anna Casey PhD [76], Dr Liz Ratcliffe PhD [76], Dr Christopher R Jones PhD [79, 105], Dr Bridget A Knight PhD [79, 105], Dr Tanzina Haque PhD, FRCPath [80], Dr Jennifer Hart MRCP [80], Dr Dianne Irish-Tavares

FRCPath [80], Eric Witele MSc [80], Craig Mower BA [86], Louisa K Watson DipHE [86], Jennifer Collins BSc [89], Gary Eltringham BSc [89], Dorian Crudgington [98], Ben Macklin [98], Prof Miren Iturriza-Gomara PhD [107], Dr Anita O Lucaci PhD [107] and Dr Patrick C McClure PhD [113]

**Metadata curation, and Sequencing and analysis:**
Matthew Carlile BSc [18], Dr Nadine Holmes PhD [18], Dr Christopher Moore PhD [18], Dr Nathaniel Storey PhD [29], Dr Stefan Rooke PhD [73], Dr Gonzalo Yebra PhD [73], Dr Noel Craine DPhil [74], Malorie Perry MSc [74], Dr Nabil-Fareed Alikhan PhD [75], Dr Stephen Bridgett PhD [77], Kate F Cook MScR [84], Christopher Fearn MSc [84], Dr Salman Goudarzi PhD [84], Prof Ronan A Lyons MD [88], Dr Thomas Williams MD [104], Dr Sam T Haldenby PhD [107], Jillian Durham BSc [116] and Dr Steven Leonard PhD [116]

**Metadata curation, and Software and analysis tools:**
Robert M Davies MA (Cantab) [116]

**Project administration, and Samples and logistics:**
Dr Rahul Batra MD [12], Beth Blane BSc [20], Dr Moira J Spyer PhD [30, 95, 96], Perminder Smith MSc [32, 112], Mehmet Yavus [85, 109], Dr Rachel J Williams PhD [96], Dr Adhyana IK Mahanama MD [97], Dr Buddhini Samaraweera MD [97], Sophia T Girgis MSc [102], Samantha E Hansford CSci [109], Dr Angie Green PhD [115], Dr Charlotte Beaver PhD [116], Katherine L Bellis [116, 102], Matthew J Dorman [116], Sally Kay [116], Liam Prestwood [116] and Dr Shavanthi Rajatileka PhD [116]

**Project administration, and Sequencing and analysis:**
Dr Joshua Quick PhD [43]

**Project administration, and Software and analysis tools:**
Radoslaw Poplawski BSc [43]

**Samples and logistics, and Sequencing and analysis:**
Dr Nicola Reynolds PhD [8], Andrew Mack MPhil [11], Dr Arthur Morriss PhD [11], Thomas Whalley BSc [11], Bindi Patel BSc [12], Dr Iliana Georgana PhD [24], Dr Myra Hosmillo PhD [24], Malte L Pinckert MPhil [24], Dr Joanne Stockton PhD [43], Dr John H Henderson PhD [65], Amy Hollis HND [65], Dr William Stanley PhD [65], Dr Wen C Yew PhD [65], Dr Richard Myers PhD [72], Dr Alicia Thornton PhD [72], Alexander Adams BSc [74], Tara Annett BSc [74], Dr Hibo Asad PhD [74], Alec Birchley MSc [74], Jason Coombes BSc [74], Johnathan M Evans MSc [74], Laia Fina [74], Bree Gatica-Wilcox MPhil [74], Lauren Gilbert [74], Lee Graham BSc [74], Jessica Hey BSc [74], Ember Hilvers MPH [74], Sophie Jones MSc [74], Hannah Jones [74], Sara Kumziene-Summerhayes MSc [74], Dr Caoimhe McKerr PhD [74], Jessica Powell BSc [74], Georgia Pugh [74], Sarah Taylor [74], Alexander J Trotter MRes [75], Charlotte A Williams BSc [96], Leanne M Kermack MSc [102], Benjamin H Foulkes MSc [109], Marta Gallis MSc [109], Hailey R Hornsby MSc [109], Stavroula F Louka MSc [109], Dr Manoj Pohare PhD [109], Paige Wolverson MSc [109], Peijun Zhang MSc [109], George MacIntyre-Cockett BSc [115], Amy Trebes MSc [115], Dr Robin J Moll PhD [116], Lynne Ferguson MSc [117], Dr Emily J Goldstein PhD [117], Dr Alasdair Maclean PhD [117] and Dr Rachael Tomb PhD [117]

**Samples and logistics, and Software and analysis tools:**
Dr Igor Starinskij MSc, MRCP [53]

**Sequencing and analysis, and Software and analysis tools:**
Laura Thomson BSc [5], Joel Southgate MSc [11, 74], Dr Moritz UG Kraemer DPhil [23], Dr Jayna Raghwani PhD [23], Dr Alex E Zarebski PhD [23], Olivia Boyd MSc [39], Lily Geidelberg MSc [39], Dr Chris J Illingworth PhD [52], Dr Chris Jackson PhD [52], Dr David Pascall PhD [52], Dr Sreenu Vattipally PhD [53], Timothy M Freeman MPhil [109], Dr Sharon N Hsu PhD [109], Dr Benjamin B Lindsey MRCP [109], Dr Keith James PhD [116], Kevin Lewis [116], Gerry Tonkin-Hill [116] and Dr Jaime M Tovar-Corona PhD [116]

**Sequencing and analysis, and Visualisation:**
MacGregor Cox MSci [20]

**Software and analysis tools, and Visualisation:**
Dr Khalil Abudahab PhD [14, 116], Mirko Menegazzo [14], Ben EW Taylor MEng [14, 116], Dr Corin A Yeats PhD [14], Afrida Mukaddas BTech [53], Derek W Wright MSc [53], Dr Leonardo de Oliveira Martins PhD [75], Dr Rachel Colquhoun DPhil [104], Verity Hill [104], Dr Ben Jackson PhD [104], Dr JT McCrone PhD [104], Dr Nathan Medd PhD [104], Dr Emily Scher PhD [104] and Jon-Paul Keatley [116]

**Leadership and supervision:**
Dr Tanya Curran PhD [3], Dr Sian Morgan FRCPath [10], Prof Patrick Maxwell PhD [20], Prof Ken Smith PhD [20], Dr Sahar Eldirdiri MBBS, MSc, FRCPath [21], Anita Kenyon MSc [21], Prof Alison H Holmes MD [38, 57], Dr James R Price PhD [38, 57], Dr Tim Wyatt PhD [69], Dr Alison E Mather PhD [75], Dr Timofey Skvortsov PhD [77] and Prof John A Hartley PhD [96]

**Metadata curation:**
Prof Martyn Guest PhD [11], Dr Christine Kitchen PhD [11], Dr Ian Merrick PhD [11], Robert Munn BSc [11], Dr Beatrice Bertolusso Degree [33], Dr Jessica Lynch MBCHB [33], Dr Gabrielle Vernet MBBS [33], Stuart Kirk MSc [34], Dr Elizabeth Wastnedge MD [56], Dr Rachael Stanley PhD [58], Giles Idle [64], Dr Declan T Bradley PhD [69, 77], Nicholas F Killough MSc [69], Dr Jennifer Poyner MD [79] and Matilde Mori BSc [110]

**Project administration:**
Owen Jones BSc [11], Victoria Wright BSc [18], Ellena Brooks MA [20], Carol M Churcher BSc [20], Dr Laia Delgado Callico PhD [20], Mireille Fragakis HND [20], Dr Katerina Galai PhD [20, 70], Dr Andrew Jermy PhD [20], Sarah Judges BA [20], Anna Markov BSc [20], Georgina M McManus BSc [20], Kim S Smith [20], Peter M D Thomas-McEwen MSc [20], Dr Elaine Westwick PhD [20], Dr Stephen W Attwood PhD [23], Dr Frances Bolt PhD [38, 57], Dr Alisha Davies PhD [74], Elen De Lacy MPH [74], Fatima Downing [74], Sue Edwards [74], Lizzie Meadows MA [75], Sarah Jeremiah MSc [97], Dr Nikki Smith PhD [109] and Luke Foulser [116]

**Samples and logistics:**
Amita Patel BSc [12], Dr Louise Berry PhD [15], Dr Tim Boswell PhD [15], Dr Vicki M Fleming PhD [15], Dr Hannah C Howson-Wells PhD [15], Dr Amelia Joseph PhD [15], Manjinder Khakh [15], Dr Michelle M Lister PhD [15], Paul W Bird MSc, MRes [16], Karlie Fallon [16], Thomas Helmer [16], Dr Claire L McMurray PhD [16], Mina Odedra BSc [16], Jessica Shaw BSc [16], Dr Julian W Tang PhD [16], Nicholas J Willford MSc [16], Victoria Blakey BSc [17], Dr Veena Raviprakash MD [17], Nicola Sheriff BSc [17], Lesley-Anne Williams BSc [17], Theresa Feltwell MSc [20], Dr Luke Bedford PhD [26], Dr James S Cargill PhD [27], Warwick Hughes MSc [27], Dr Jonathan Moore

MD [28], Susanne Stonehouse BSc [28], Laura Atkinson MSc [29], Jack CD Lee MSc [29], Dr Divya Shah PhD [29], Natasha Ohemeng-Kumi MSc [32, 112], John Ramble MSc [32, 112], Jasveen Sehmi MSc [32, 112], Dr Rebecca Williams BMBS [33], Wendy Chatterton MSc [34], Monika Pusok MSc [34], William Everson MSc [37], Anibolina Castigador IBMS HCPC [44], Emily Macnaughton FRCPath [44], Dr Kate El Bouzidi MRCP [45], Dr Temi Lampejo FRCPath [45], Dr Malur Sudhanva FRCPath [45], Cassie Breen BSc [47], Dr Graciela Sluga MD, MSc [48], Dr Shazaad SY Ahmad MSc [49, 70], Dr Ryan P George PhD [49], Dr Nicholas W Machin MSc [49, 70], Debbie Binns BSc [50], Victoria James BSc [50], Dr Rachel Blacow MBCHB [55], Dr Lindsay Coupland PhD [58], Dr Louise Smith PhD [59], Dr Edward Barton MD [60], Debra Padgett BSc [60], Garren Scott BSc [60], Dr Aidan Cross MBCHB [61], Dr Mariyam Mirfenderesky FRCPath [61], Jane Greenaway MSc [62], Kevin Cole [64], Phillip Clarke [67], Nichola Duckworth [67], Sarah Walsh [67], Kelly Bicknell [68], Robert Impey MSc [68], Dr Sarah Wyllie PhD [68], Richard Hopes [70], Dr Chloe Bishop PhD [72], Dr Vicki Chalker PhD [72], Dr Ian Harrison PhD [72], Laura Gifford MSc [74], Dr Zoltan Molnar PhD [77], Dr Cressida Auckland FRCPath [79], Dr Cariad Evans PhD [85, 109], Dr Kate Johnson PhD [85, 109], Dr David G Partridge FRCP, FRCPath [85, 109], Dr Mohammad Raza PhD [85, 109], Paul Baker MD [86], Prof Stephen Bonner PhD [86], Sarah Essex [86], Leanne J Murray [86], Andrew I Lawton MSc [87], Dr Shirelle Burton-Fanning MD [89], Dr Brendan AI Payne MD [89], Dr Sheila Waugh MD [89], Andrea N Gomes MSc [91], Maimuna Kimuli MSc [91], Darren R Murray MSc [91], Paula Ashfield MSc [92], Dr Donald Dobie MBCHB [92], Dr Fiona Ashford PhD [93], Dr Angus Best PhD [93], Dr Liam Crawford PhD [93], Dr Nicola Cumley PhD [93], Dr Megan Mayhew PhD [93], Dr Oliver Megram PhD [93], Dr Jeremy Mirza PhD [93], Dr Emma Moles-Garcia PhD [93], Dr Benita Percival PhD [93], Megan Driscoll BSc [96], Leah Ensell BSc [96], Dr Helen L Lowe PhD [96], Laurentiu Maftei BSc [96], Matteo Mondani MSc [96], Nicola J Chaloner BSc [99], Benjamin J Cogger BSc [99], Lisa J Easton MSc [99], Hannah Huckson BSc [99], Jonathan Lewis MSc, PgD, FIBMS [99], Sarah Lowdon BSc [99], Cassandra S Malone MSc [99], Florence Munemo BSc [99], Manasa Mutingwende MSc [99], Roberto Nicodemi BSc [99], Olga Podplomyk FD [99], Thomas Somassa BSc [99], Dr Andrew Beggs PhD [100], Dr Alex Richter PhD [100], Claire Cormie [102], Joana Dias MSc [102], Sally Forrest BSc [102], Dr Ellen E Higginson PhD [102], Mailis Maes MPhil [102], Jamie Young BSc [102], Dr Rose K Davidson PhD [103], Kathryn A Jackson MSc [107], Dr Alexander J Keeley MRCP [109], Prof Jonathan Ball PhD [113], Timothy Byaruhanga MSc [113], Dr Joseph G Chappell PhD [113], Jayasree Dey MSc [113], Jack D Hill MSc [113], Emily J Park MSc [113], Arezou Fanaie MSc [114], Rachel A Hilson MSc [114], Geraldine Yaze MSc [114] and Stephanie Lo [116]

**Sequencing and analysis:**

Safiah Afifi BSc [10], Robert Beer BSc [10], Joshua Maksimovic FD [10], Kathryn McCluggage Masters [10], Karla Spellman FD [10], Catherine Bresner BSc [11], William Fuller BSc [11], Dr Angela Marchbank BSc [11], Trudy Workman HNC [11], Dr Ekaterina Shelest PhD [13, 81], Dr Johnny Debebe PhD [18], Dr Fei Sang PhD [18], Dr Sarah Francois PhD [23], Bernardo Gutierrez MSc [23], Dr Tetyana I Vasylyeva DPhil [23], Dr Flavia Flaviani PhD [31], Dr Manon Ragonnet-Cronin PhD [39], Dr Katherine L Smollett PhD [42], Alice Broos BSc [53], Daniel Mair BSc [53], Jenna Nichols BSc [53], Dr Kyriaki Nomikou PhD [53], Dr Lily Tong PhD [53], Ioulia Tsatsani MSc [53], Prof Sarah O'Brien PhD [54], Prof Steven Rushton PhD [54], Dr Roy Sanderson PhD [54], Dr Jon Perkins MBCHB [55], Seb Cotton MSc [56], Abbie Gallagher BSc [56], Dr Elias Allara MD, PhD [70, 102], Clare Pearson MSc [70, 102], Dr David Bibby PhD [72], Dr Gavin Dabrera PhD [72], Dr Nicholas Ellaby PhD [72], Dr Eileen Gallagher PhD [72], Dr Jonathan Hubb PhD [72], Dr Angie Lackenby PhD [72], Dr David Lee PhD [72], Nikos Manesis [72], Dr Tamyo Mbisa PhD [72], Dr Steven Platt PhD [72], Katherine A Twohig [72], Dr Mari Morgan PhD [74], Alp Aydin MSci [75], David J Baker BEng [75], Dr Ebenezer Foster-Nyarko PhD [75], Dr Sophie J Prosolek PhD [75], Steven Rudder [75], Chris

Baxter BSc [77], Sílvia F Carvalho MSc [77], Dr Deborah Lavin PhD [77], Dr Arun Mariappan PhD [77], Dr Clara Radulescu PhD [77], Dr Aditi Singh PhD [77], Miao Tang MD [77], Helen Morcrette BSc [79], Nadua Bayzid BSc [96], Marius Cotic MSc [96], Dr Carlos E Balcazar PhD [104], Dr Michael D Gallagher PhD [104], Dr Daniel Maloney PhD [104], Thomas D Stanton BSc [104], Dr Kathleen A Williamson PhD [104], Dr Robin Manley PhD [105], Michelle L Michelsen BSc [105], Dr Christine M Sambles PhD [105], Dr David J Studholme PhD [105], Joanna Warwick-Dugdale BSc [105], Richard Eccles MSc [107], Matthew Gemmell MSc [107], Dr Richard Gregory PhD [107], Dr Margaret Hughes PhD [107], Charlotte Nelson MSc [107], Dr Lucille Rainbow PhD [107], Dr Edith E Vamos PhD [107], Hermione J Webster BSc [107], Dr Mark Whitehead PhD [107], Claudia Wierzbicki BSc [107], Dr Adrienn Angyal PhD [109], Dr Luke R Green PhD [109], Dr Max Whiteley PhD [109], Emma Betteridge BSc [116], Dr Iraad F Bronner PhD [116], Ben W Farr BSc [116], Scott Goodwin MSc [116], Dr Stefanie V Lensing PhD [116], Shane A McCarthy [116, 102], Dr Michael A Quail PhD [116], Diana Rajan MSc [116], Dr Nicholas M Redshaw PhD [116], Carol Scott [116], Lesley Shirley MSc [116] and Scott AJ Thurston BSc [116]

**Software and analysis tools:**
Dr Will Rowe PhD[43], Amy Gaskin MSc [74], Dr Thanh Le-Viet PhD [75], James Bonfield BSc [116], Jennifier Liddle [116] and Andrew Whitwham BSc [116]

**1** Barking, Havering and Redbridge University Hospitals NHS Trust, **2** Barts Health NHS Trust, **3** Belfast Health & Social Care Trust, **4** Betsi Cadwaladr University Health Board, **5** Big Data Institute, Nuffield Department of Medicine, University of Oxford, **6** Blackpool Teaching Hospitals NHS Foundation Trust, **7** Bournemouth University, **8** Cambridge Stem Cell Institute, University of Cambridge, **9** Cambridge University Hospitals NHS Foundation Trust, **10** Cardiff and Vale University Health Board, **11** Cardiff University, **12** Centre for Clinical Infection and Diagnostics Research, Department of Infectious Diseases, Guy's and St Thomas' NHS Foundation Trust, **13** Centre for Enzyme Innovation, University of Portsmouth, **14** Centre for Genomic Pathogen Surveillance, University of Oxford, **15** Clinical Microbiology Department, Queens Medical Centre, Nottingham University Hospitals NHS Trust, **16** Clinical Microbiology, University Hospitals of Leicester NHS Trust, **17** County Durham and Darlington NHS Foundation Trust, **18** Deep Seq, School of Life Sciences, Queens Medical Centre, University of Nottingham, **19** Department of Infectious Diseases and Microbiology, Cambridge University Hospitals NHS Foundation Trust, **20** Department of Medicine, University of Cambridge, **21** Department of Microbiology, Kettering General Hospital, **22** Department of Microbiology, South West London Pathology, **23** Department of Zoology, University of Oxford, **24** Division of Virology, Department of Pathology, University of Cambridge, **25** East Kent Hospitals University NHS Foundation Trust, **26** East Suffolk and North Essex NHS Foundation Trust, **27** East Sussex Healthcare NHS Trust**,** **28** Gateshead Health NHS Foundation Trust, **29** Great Ormond Street Hospital for Children NHS Foundation Trust, **30** Great Ormond Street Institute of Child Health (GOS ICH), University College London (UCL), **31** Guy's and St. Thomas' Biomedical Research Centre, **32** Guy's and St. Thomas' NHS Foundation Trust, **33** Hampshire Hospitals NHS Foundation Trust, **34** Health Services Laboratories, **35** Heartlands Hospital, Birmingham, **36** Hub for Biotechnology in the Built Environment, Northumbria University, **37** Hull University Teaching Hospitals NHS Trust, **38** Imperial College Healthcare NHS Trust, **39** Imperial College London, **40** Infection Care Group, St George's University Hospitals NHS Foundation Trust, **41** Institute for Infection and Immunity, St George's University of London, **42** Institute of Biodiversity, Animal Health & Comparative Medicine, **43** Institute of Microbiology and Infection, University of Birmingham, **44** Isle of Wight NHS Trust, **45** King's College Hospital NHS Foundation Trust, **46** King's College London, **47** Liverpool Clinical Laboratories, **48** Maidstone and Tunbridge Wells NHS Trust, **49** Manchester University NHS Foundation Trust, **50** Microbiology Department, Buckinghamshire Healthcare NHS Trust, **51** Microbiology, Royal Oldham Hospital, **52** MRC Biostatistics Unit, University of Cambridge, **53** MRC-

University of Glasgow Centre for Virus Research, **54** Newcastle University, **55** NHS Greater Glasgow and Clyde, **56** NHS Lothian, **57** NIHR Health Protection Research Unit in HCAI and AMR, Imperial College London, **58** Norfolk and Norwich University Hospitals NHS Foundation Trust, **59** Norfolk County Council, **60** North Cumbria Integrated Care NHS Foundation Trust, **61** North Middlesex University Hospital NHS Trust, **62** North Tees and Hartlepool NHS Foundation Trust, **63** North West London Pathology, **64** Northumbria Healthcare NHS Foundation Trust, **65** Northumbria University, **66** NU-OMICS, Northumbria University, **67** Path Links, Northern Lincolnshire and Goole NHS Foundation Trust, **68** Portsmouth Hospitals University NHS Trust, **69** Public Health Agency, Northern Ireland, **70** Public Health England, **71** Public Health England, Cambridge, **72** Public Health England, Colindale, **73** Public Health Scotland, **74** Public Health Wales, **75** Quadram Institute Bioscience, **76** Queen Elizabeth Hospital, Birmingham, **77** Queen's University Belfast, **78** Royal Brompton and Harefield Hospitals, **79** Royal Devon and Exeter NHS Foundation Trust, **80** Royal Free London NHS Foundation Trust, **81** School of Biological Sciences, University of Portsmouth, **82** School of Health Sciences, University of Southampton, **83** School of Medicine, University of Southampton, **84** School of Pharmacy & Biomedical Sciences, University of Portsmouth, **85** Sheffield Teaching Hospitals NHS Foundation Trust, **86** South Tees Hospitals NHS Foundation Trust, **87** Southwest Pathology Services, **88** Swansea University, **89** The Newcastle upon Tyne Hospitals NHS Foundation Trust, **90** The Queen Elizabeth Hospital King's Lynn NHS Foundation Trust, **91** The Royal Marsden NHS Foundation Trust, **92** The Royal Wolverhampton NHS Trust, **93** Turnkey Laboratory, University of Birmingham, **94** University College London Division of Infection and Immunity**,** **95** University College London Hospital Advanced Pathogen Diagnostics Unit**,** **96** University College London Hospitals NHS Foundation Trust, **97** University Hospital Southampton NHS Foundation Trust, **98** University Hospitals Dorset NHS Foundation Trust, **99** University Hospitals Sussex NHS Foundation Trust, **100** University of Birmingham, **101** University of Brighton, **102** University of Cambridge, **103** University of East Anglia, **104** University of Edinburgh, **105** University of Exeter, **106** University of Kent, **107** University of Liverpool, **108** University of Oxford, **109** University of Sheffield, **110** University of Southampton, **111** University of St Andrews, **112** Viapath, Guy's and St Thomas' NHS Foundation Trust, and King's College Hospital NHS Foundation Trust, **113** Virology, School of Life Sciences, Queens Medical Centre, University of Nottingham, **114** Watford General Hospital, **115** Wellcome Centre for Human Genetics, Nuffield Department of Medicine, University of Oxford, **116** Wellcome Sanger Institute, **117** West of Scotland Specialist Virology Centre, NHS Greater Glasgow and Clyde, **118** Whittington Health NHS Trust
